# Genetic Heterogeneity Profiling by Single Cell RNA Sequencing

**DOI:** 10.1101/457622

**Authors:** Zilu Zhou, Bihui Xu, Andy Minn, Nancy R Zhang

## Abstract

Detection of genetically distinct subclones and profiling the transcriptomic differences between them is important for studying the evolutionary dynamics of tumors, as well as for accurate prognosis and effective treatment of cancer in the clinic. For the profiling of intra-tumor transcriptional heterogeneity, single cell RNA-sequencing (scRNA-seq) is now ubiquitously adopted in ongoing and planned cancer studies. Detection of somatic DNA mutations and inference of clonal membership from scRNA-seq, however, is currently unreliable. We propose DENDRO, an analysis method for scRNA-seq data that detects genetically distinct subclones, assigns each single cell to a subclone, and reconstructs the phylogenetic tree describing the tumor’s evolutionary history. DENDRO utilizes information from single nucleotide mutations in transcribed regions and accounts for technical noise and expression stochasticity at the single cell level. The accuracy of DENDRO was benchmarked on spike-in datasets and on scRNA-seq data with known subpopulation structure. We applied DENDRO to delineate subclonal expansion in a mouse melanoma model in response to immunotherapy, highlighting the role of neoantigens in treatment response. We also applied DENDRO to primary and lymph-node metastasis samples in breast cancer, where the new approach allowed us to better understand the relationship between genetic and transcriptomic intratumor variation.

## Background

DNA alterations, especially single nucleotide alteration (SNA) and epigenetic modulation both contribute to intratumor heterogeneity [1], which mediates tumor initiation, progression, metastasis and relapse [2, 3]. Intratumor genetic and transcriptomic variation underlie patients’ response to treatment, as natural selection can lead to the emergence of subclones that are drug resistant [4]. Thus, identifying subclonal DNA alterations and assessing their impact on intratumor transcriptional dynamics can elucidate the mechanisms of tumor evolution and, further, uncover potential targets for therapy. To characterize intratumor genetic heterogeneity, most prior studies have used bulk tumor DNA sequencing [5-12], but these approaches have limited resolution and power [13].

Breakthroughs in single-cell genomics promise to reshape cancer research by allowing comprehensive cell type classification and rare subclone identification. For example, in breast cancer, single-cell DNA sequencing (scDNA-seq) was used to distinguish normal cells from malignant cells, the latter of which were further classified into subclones [14-16]. For the profiling of intra-tumor transcriptional heterogeneity, single cell RNA-sequencing (scRNA-seq), such as Smart-seq2 [17], Drop-seq [18], and 10X Genomics Chromium™, is now ubiquitously adopted in ongoing and planned cancer studies. ScRNA-seq studies have already led to novel insights into cancer progression and metastasis, as well as into tumor prognosis and treatment response, especially response variability in immune checkpoint blockade (ICB) [19-26]. Characterization of intratumor genetic heterogeneity and identification of subclones using scRNA-seq is challenging, as SNAs derived from scRNA-seq reads are extremely noisy and most studies have relied on the detection of chromosome-level copy number aberrations through smoothed gene expression profiles. Yet, as intratumor transcriptomic variation is partially driven by intratumor genetic variation, the classification of cells into subclones and the characterization of each subclone’s genetic alterations should ideally be an integral step in any scRNA-seq analysis.

The appeal of subclone identification in scRNA-seq data is compounded by the shortage of technology for sequencing the DNA and RNA molecules in the *same* cell with acceptable accuracy, throughput, and cost. Although one can apply both scDNA-seq and scRNA-seq to a given cell population, the mutation analysis and RNA quantification cannot be conducted in the same set of cells. Although there are now technologies for deep targeted sequencing of select transcripts matched with same-cell whole transcriptome sequencing [27, 28], these methods are still, in effect, profiling DNA-level variation by sequencing expressed transcripts, and are thus subject to the technical issues, especially dropout due to transcriptional stochasticity, that we modeled in DENDRO.

Subclone detection using scRNA-seq is difficult mainly because only a small portion of the SNAs of each cell is expected to be seen in the read output of scRNA-seq. This is because to be sequenced, an SNA needs to fall in a transcribed region of the genome, at a location within the transcript that will eventually be read by the chosen sequencing protocol. Even for SNAs that satisfy these requirements, the mutated allele are often missing in the read output due to *dropout*, especially in the heterozygous case. This is due, in part, to the bursty nature of gene transcription in single cells [29-31], where in any given cell, a substantial fraction of the genes are only expressed from one of the alleles. Thus, an SNA residing in a gene that is expressed at the bulk tissue level may not be observed in a particular cell, simply because the mutated allele, by chance, is not expressed in the given cell. We refer to alleles that are not captured due to expression stochasticity as *biological dropouts*. Even for a mutated allele that is expressed, it has to be successfully converted to cDNA and then sequenced to be represented in the final read output; we refer to alleles lost due to technical reasons as *technical dropouts*. In addition to dropout events, post-transcriptional modification, such as RNA editing, and sequencing errors impede both the sensitivity and the specificity of SNA discovery. As a result, methods developed for single cell SNA detection using scDNA-seq, such as Monovar [32], as well as methods designed for SNA detection in bulk DNA or RNA sequencing data do not yield accurate results in the scRNA-seq setting [33-38].

Here we present a new statistical and computational framework – <u>D</u>NA based <u>E</u>volution<u>N</u>ary tree pre<u>D</u>iction by sc<u>R</u>NA-seq techn<u>O</u>logy (DENDRO) - that reconstructs the phylogenetic tree for cells sequenced by scRNA-seq based on genetic divergence calculated from DNA-level mutations. DENDRO assigns each cell to a leaf in the tree representing a subclone, and, for each subclone, infers its mutation profile. DENDRO can detect genetically divergent subclones by addressing challenges unique to scRNA-seq, including transcriptional variation and technical noise. A DENDRO clustering of scRNA-seq data allows joint genetic and transcriptomic analysis on the same set of cells.

We evaluate DENDRO against existing approaches, through spike-in data sets and a metastasized renal cell carcinoma dataset with known subpopulation labels, and show that DENDRO improved the accuracy of subclone detection. We then demonstrate the DENDRO to biological discovery through two applications. The first application profiles the treatment response in a melanoma model to immune checkpoint blockade therapy. DENDRO identified a subclone that contracted consistently in response to ICB therapy, and revealed that the contraction was driven by the high mutation burden and increased availability of predicted neoantigens. Transcriptional divergence between the subclones in this model was very weak, and thus the neoantigen-driven sub-clonal dynamics would not have been detected without extracting DNA-level information. In the second application to a breast tumor dataset, DENDRO detected subclones and allowed for the joint characterization of transcriptomic and genetic divergence between cells in lymph-node metastasis and cells in primary resections.

The DENDRO package, implemented in R, is available at https://github.com/zhouzilu/DENDRO, where we also provide a power calculation toolkit, DENDROplan, to aid in the design of scRNA-seq experiments for subclonal mutation analysis using DENDRO.

## Results

### Method overview

#### Overview of DENDRO model and pipeline

Figure 1A shows an overview of DENDRO’s analysis pipeline. Per cell counts of total read coverage (*N* matrix) and mutation allele read coverage (*X* matrix) at SNA locations are extracted after read alignment and SNA detection (details in Methods, Figure S1). Based on these matrices, DENDRO then computes a cell-to-cell genetic divergence matrix, where entry (*c, c*’) of the matrix is a measure of the genetic divergence between cells *c* and *c*’). Details of this genetic divergence evaluation will be given in the next section. DENDRO then clusters the cells into genetically distinct subclones based on this pairwise divergence matrix, and selects the number of subclones based on inspection of the intra-cluster divergence curve. Reads from the same subclone are then pooled together, and the SNA profile for each subclone is re-estimated based on the pooled reads, which improves upon the previous SNA profiles computed at the single cell level. Finally, DENDRO generates a parsimony tree using the subclone-level mutation profiles to more accurately reflect the evolutionary relationship between the subclones.

**Figure 1.**
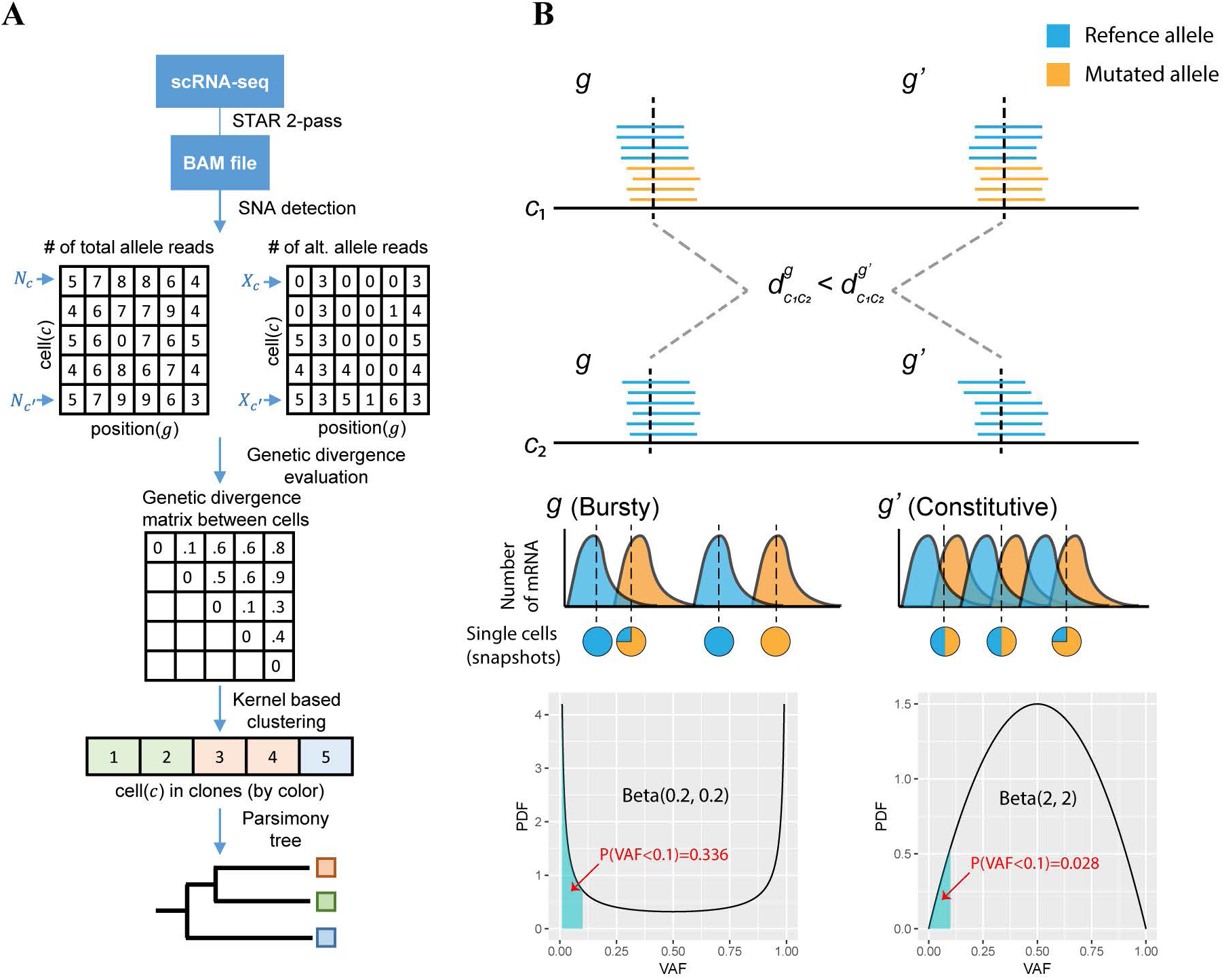
DENDRO analysis pipeline and genetic divergence evaluation. **a** DENDRO analysis pipeline overview. **b** Statistical model for genetic divergence evaluation function. In this example, we show relationship of cell *c*_*1*_ and *c*_*2*_ with gene *g* and *g’* in terms of their genetic divergence. Middle panels shows the stochasticity of cell expression and bottom panels illustrates the distribution of variants allele frequency across cells for these two genes, indicating that gene *g* is a bursty gene and *g’* is a constitutive gene.

#### Genetic divergence evaluation

Due to the high rates of biological and technical dropout, SNA detection within each individual cell lacks sensitivity. We also expect low specificity due to the high base error rate in scRNA-seq protocols. Thus, simple distance measures such as the Hamming or Euclidean distances evaluated on the raw SNA genotype matrix or the raw allele frequency matrix do not accurately reflect the genetic divergence between cells.

To more accurately estimate the cell-to-cell genetic divergence, we have developed a statistical model that accounts for technical dropout, sequencing error and expression stochasticity. Consider two cells, *c* and *c*’, and let *I*_*c*_ and *I*_*c*′_ index the clonal group to which the cells belong. That is, *I*_*c*_ = *I*_*c*′_ if cells *c* and *c*’ come from the same subclone and thus share the same SNA profile. Let *X*_*c*_ = (*X*_*c*1_,…,*X*_*cm*_) be the mutation allele read counts for this cell at the *m* SNA sites profiled, and *N*_*c*_ = (*N*_*c*1_,…,*N*_*cm*_) be the total read counts at these sites. We define the genetic divergence between the two cells as

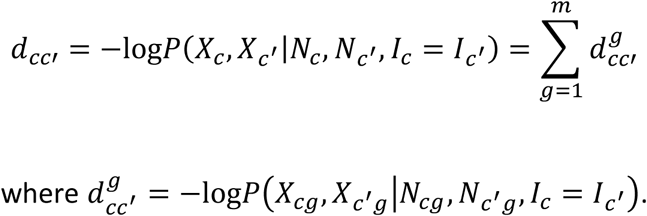

In other words, *d*_*cc*′_ is the negative log likelihood of the mutation allele counts of cells *c* and *c*′, given the total read counts and the event that the two cells belong to the same subclone. If *c* and *c*′ have mutations in mismatched positions, this likelihood for *X*_*c*_, *X*_*c*′_ conditioned on *I*_*c*_ = *I*_*c*′_ would be small, giving a large value for *d*_*cc*′_. By the assumption of independence between sites, *d*_*cc*′_ is the sum of 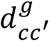 where 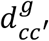 is the contribution of mutation site *g* to the divergence measure. In characterizing the conditional distribution for *X*_*cg*_ and *X*_*c*′*g*_, we use a Beta-Binomial distribution to model expression stochasticity and a Binomial model to capture sequencing errors and rare RNA-editing events. Referring to Figure 1b, mutations residing in bursty genes, such as gene *g*, would tend to have U-shaped allele frequency distributions and are more likely to be “dropped” due to low or zero expression. In contrary, mutations residing in constitutive (non-bursty) genes, such as gene *g*′ in Figure 1b, would have bell-shaped allele frequency distributions and can be genotyped more reliably. Thus, even if the read counts for the mutation loci residing in genes *g* and *g*′ are identical across two cells (*c*_1_ and *c*_2_ in Figure 1b), the locus in *g*′ would contribute a higher value, compared to the locus in *g*, to the divergence between cells *c*_1_ and *c*_2_. Please see Methods for details.

### Accuracy assessment

#### Accuracy assessment with in silico spike-in experiment

First, we designed a spike-in procedure to assess the accuracy of DENDRO versus existing approaches and to make realistic power projections for subclone detection (Figure 2a). Since DENDRO is currently the only method for SNA-based subclone detection using scRNA-seq data alone, we benchmarked against more straightforward approaches such as hierarchical clustering based on mutation allele frequencies. The spike-in procedure starts with an assumed evolutionary tree, where the leaves are subclones and mutations can be placed on the branches. In the absence of prior information, a simple tree structure is used, such as the one shown in Figure 2A. Parameters of simulation are (1) total number of mutations, (2) total number of cells, (3) the proportion of cells in each clade, (4) the proportion of mutations along each branch, and (5) mean read coverage across loci. Some of these parameters can be determined using bulk DNA-seq and/or bulk RNA-seq data if available (Methods). Parameters (1-4) determine the mutation profile matrix *Z*. To get the matrix of alternative and total read counts for each mutation loci in each cell (*X*_*cg*_ and *N*_*cg*_, respectively), we overlay the mutation matrix onto the scRNA-seq data. This allows the spike-in framework to retain the expression stochasticity and sequencing error of real scRNA-seq data. DENDRO is then applied to the read count matrices to obtain the subclone clusters, which is then compared with the known labels. Accuracy is evaluated by three metrics: adjusted Rand index, capture rate and purity (Supplementary Materials). Such spike-in procedure can also facilitate experiment design, as it predicts the expected clustering accuracy by DENDRO given sequencing parameters and available bulk data for the tumor (See DENDROplan in Methods).

**Figure 2.**
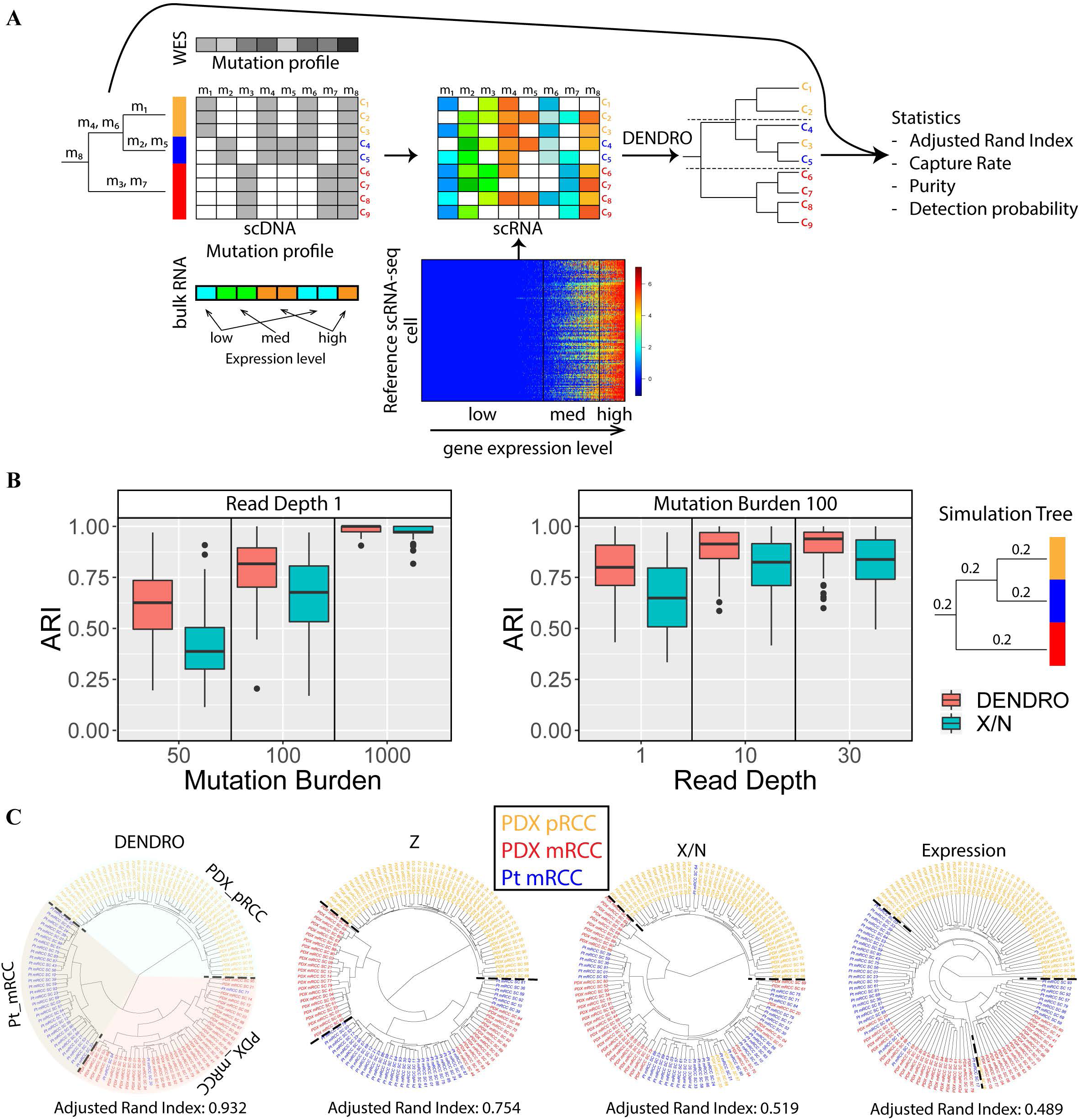
DENDRO accuracy assessment. **a** The overall spike-in analysis pipeline. Three statistics measure the accuracy of DENDRO - adjusted Rand index (global accuracy, 1 indicating perfect classification), capture rate (subclone specific accuracy, out of all cells from the clade, how many of them are correctly assigned) and purity (subclone specific accuracy, out of all cells from the detected cluster, how many are actually from the clade). **b** Cluster accuracy via spike-in studies. Various parameters show effects on cluster accuracy (measured by adjusted Rand index) based on tree structure on the right. Left panel: effect of mutation read depth. Right panel: effect of total mutation burden. **c** Evaluation of DENDRO on a renal cell carcinoma and its metastasis. (Left to right) (1) DENDRO clustering result from primary and metastatic renal cell carcinoma dataset. Background colors represent DEDRO clustering result. (2) Clustering of the same dataset using *Z* matrix (indicator matrix, *z*_*ij*_ = 1 when detected a mutation for cell i at site *j* by GATK tool). (3) Clustering of the same dataset using *X/N* matrix (mutation allele frequency matrix) (4) Clustering of the same dataset using expression (*logT-*PM).

Using the above framework, we conducted a systematic evaluation of DENDRO’s subclone detection accuracy on an example scRNA-seq dataset with allelic information [39]. The results, compiled in Figure 2b, Supplementary Materials and Figure S3, show that DENDRO has better performance than simply clustering on mutation allele frequencies and quantify how accuracy depends on the mutation burden, mutation read depth, mutation distribution, subclone cell proportion, and cell populations. Even when there are only 100 mutations with relatively low average coverage (read depth equals to 1), DENDRO can still extract meaningful clustering results (average ARI ≈ 0.8). More importantly, variation in total expression of genes does not influence DENDRO’s divergence measure. DENDRO shows consistent results in spike-in analysis between populations of single cell type and multiple cell types (Figure S3). This is due to DENDRO’s reliance only on the distribution of the mutation allele frequency conditioned on the total read coverage, as illustrated by the simulation study (Supplementary Material and Figure S2). The divergence evaluation reflects solely genetic distance not transcriptomic difference, allowing for easy interpretation. A more extensive spike-in analysis can be found in the Supplementary Materials.

#### Accuracy assessment on a renal cell carcinoma and its metastasis

We also benchmarked DENDRO against existing methods on the renal cell carcinoma dataset from Kim et al [21] (Figure 2C). This dataset contained 116 cells sequenced using the Smart-seq technology [17], obtained from three tumors derived from one patient: a patient-derived xenograft (PDX) from the primary renal cell carcinoma (PDX_pRCC), a biopsy of the metastasis to the lung 1 year after treatment of primary site (Pt_mRCC), and a PDX of the lung metastasis renal cell carcinoma (PDX_mRCC) (Figure S4A). The cells should share common early driver mutations due to their shared origin from the same patient, but the metastasis and the cultivation of each tumor in separate medium (human or mouse) should have allowed for the accumulation of new mutations. Thus, we expect the three tumors to be clonally distinct. This knowledge allows us to use this dataset to benchmark accuracy and to illustrate how DENDRO enables joint analysis of the genetic and transcriptomic heterogeneity at single cell resolution.

We compared 4 different clustering methods: (1) DENDRO, (2) hierarchical clustering based on the primary genotype matrix *Z* generated by GATK (*Z*_*cg*_ = 1 when a mutation *g* is detected for cell *c, Z*_*cg*_ = 0 otherwise), (3) hierarchical clustering based on the *X*/*N* matrix that preserve the variant allele information and (4) hierarchical clustering based on gene expression (log *TPM*). DENDRO gives the cleanest separation between the three populations with adjusted Rand Index of 0.932 (1.0 indicates perfect clustering, Figure 2C panel 1), as compared to 0.754 for Z matrix (Figure 2C panel 2), 0.519 for X/N (Figure 2C panel 3) and 0.489 for expression (Figure 2C panel 4). Inspection of the tree shows that, as expected, divergence between primary tumor and metastasis exceeds divergence between patient sample and PDX sample, as PDX_mRCC clusters with Pt_mRCC rather than PDX_pRCC. All of the other three methods successfully separated the primary sample from the metastatic samples, but could not differentiate between the two metastasis samples.

For DENDRO, the intra-cluster divergence curve flattened at 3, and thus we stopped splitting at 3 clusters (Figure S4E and Methods). We annotated the clusters as PDX_mRCC, PDX_pRCC and Pt_mRCC by their cell compositions (Table S3A). DENDRO found minimal sharing of subclones among the tumors derived from three sources, and low genetic heterogeneity within each tumor. This is unsurprising since relapsed metastasis consists of cells that have already undergone selection, and since the PDX tumors are each seeded by a small subsample of cells from the original tumor, each tumor consists of unique subclones not detected in other sites [40-42]. Additional joint analysis of transcriptome and DNA mutations can be found in the Supplementary Materials.

### DENDRO analysis of melanoma model in response to immune checkpoint blockade highlights the role of neoantigens

Immune checkpoint blockade (ICB) of the inhibitory receptors CTLA4 and PD1 can result in durable responses in multiple cancer types [43]. Features intrinsic to cancer cells that can impact ICB treatment outcome include their repertoire of neoantigens [44], tumor mutational burden (TMB) [45], and expression of PDL1 [46]. DENDRO analysis of scRNA-seq data allows joint DNA-RNA analysis of single cells, thus enabling the simultaneous quantification of tumor mutational burden, the prediction of neoantigen repertoire, and the characterization of gene expression profile at subclonal resolution. Thus, to demonstrate the power of DENDRO and to better understand the relationship between ICB response and intratumor heterogeneity, we profiled the single cell transcriptomes across three conditions derived from 2 melanoma cell lines (Figure 3A): B16 melanoma cell line, which has shown modest initial response to ICB treatment but eventually grows out, and Res 499 melanoma cell line (R499), which was derived from a relapsed B16 tumor after combined treatment of radiation and anti-CTLA4 and is fully resistant to ICB [47]. B16 was evaluated with and without anti-PD1 treatment, as we wanted a tumor model that captures a transient ICB response. A total of 600 tumor cells were sequenced with Smart-seq technology from six mice across three conditions: two mice with B16 without treatment (B16), two mice with B16 after anti-PD1 treatment (B16PD1) and two mice with R499 without treatment (R499) (Figure 3A and Methods). The existence of multiple subclones in B16 and R499 was suggested by bulk WES analysis [47, 48]. Our goal here is to determine whether the subclones differ in anti-PD1 response, and if so, what are the subclonal differences.

**Figure 3.**
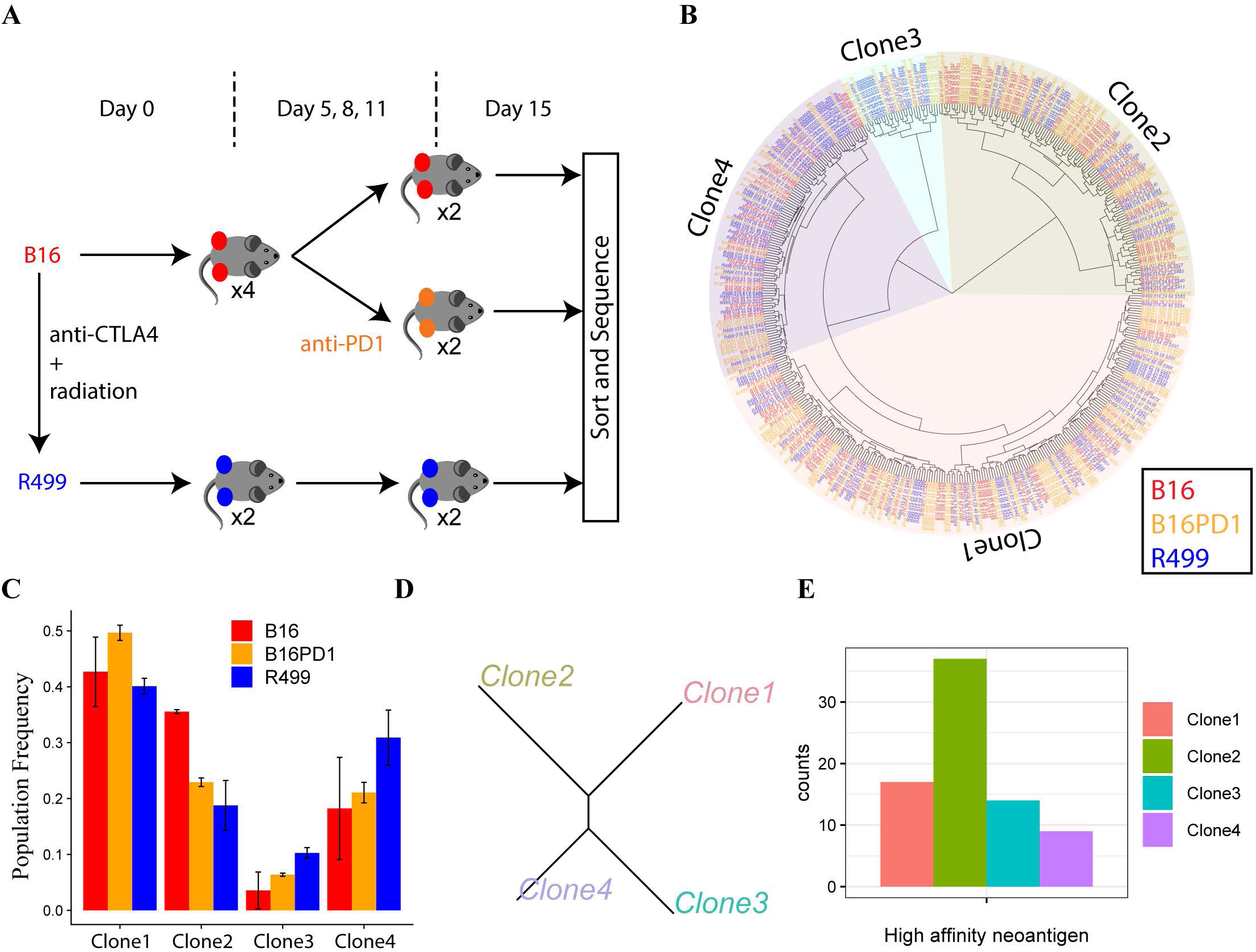
Clonal composition alternations with anti-PDl treatments and cell lines. **a** Experimental overview. For each condition at Day 15, we have two biological replicates. There are total 600 cells from 6 tumors sequenced. **b** DENDRO cluster result. No clone is exclusively associated with any tumor condition. **c** Frequencies of the subclonal populations in B16, B16PD1 and R499. **d** Neighbor joining phylogenetic tree given detected subclones. **e** Number of high affinity neoantigen predicted for each clone. Clone 2 have the highest number of neoantigen.

A DENDRO analysis of 4059 putative mutation sites across 460 cells retained after QC (see Methods and Figure S9A, B, C) yields the clustering displayed in Figure 3B, with four subclones suggested by the intracluster divergence curve (Figure S9D). All subclones are shared among the three conditions, which is not unexpected given that all tumor cells were derived from the same parental cell line. However, the subclonal proportions vary significantly between conditions (Figure 3B). The subclonal proportions of B16PD1 are approximately intermediate between that of B16 and R499. This is expected as R499 had gone through immune editing whereas B16PD1, at the time of harvest, was still undergoing immune editing and was at the transient response state. Furthermore, the selective pressure of radiation plus anti-CTLA4 is likely more than that of anti-PD1 treatment, as the former but not the latter results in complete responses in our B16 model [47]. The frequency of Clone 2 is lower in B16PD1 and R499, indicating sensitivity to anti-PD1 treatment, while the frequencies of Clone 3 and Clone 4 increase after treatment and are the highest in R499, indicating resistance to therapy (Figure 3C, S10A).

To explore why subclones vary in sensitivity to anti-PD1 treatment, we compared the mutation profile of Clone 2 to the other subclones. We pooled cells in each of the four subclones and re-estimated their mutation profiles, which were then used to construct a phylogenetic tree (Figure 3D). The phylogeny suggests that Clone 3 and Clone 4 are genetically closer to each other than to Clone 2, and thus, their similarity in treatment response may be in part due to similarity in their mutation profiles. The re-estimated mutation profiles show that Clone 2 has the highest tumor mutation burden, which has been associated with increased likelihood of ICB response [49, 50]. We then predicted the quantity of high-affinity (≤ 100 nm) neoantigens in each subclone given its mutation profile [48]. As shown in Figure 3E, Clone 2 has twice as many high-affinity neoantigens as the other three subclones. The high level of neoantigens can lead to better T cell recognition, resulting in increased efficacy of anti-PD1 treatment 252 [51].

Analysis of gene expression, on the other hand, did not yield detectable known signatures associated with anti-PD1 treatment sensitivity. Projections based on the expression of highly variable genes, as shown in PCA and t-SNE plots (Figure S8), did not yield meaningful clusters. Differential expression analysis between each subclone and the other subclones found few genes with adjusted P-value < 0.05, indicating similar expression across sub-clones that is concordant with the lack of structure in the expression PCA and tSNE plots. Expressions of *Pdl1* (aka. *Cd274*) showed no differences between subclones (KS-test: P-value > 0.42, Figure S10B). In addition, there were no detectable chromosome-level differences in smoothed gene expression, indicating that there are no large CNV events that distinguish the subclones (Figure S11). DENDRO, detecting exonic mutations from scRNA-seq data, enabled the finding of subclones in this data, the prediction of neoantigen load of each subclone, and the analysis of subclonal dynamics due to treatment. Our analysis suggests that the genetic heterogeneity, rather than transcriptomic heterogeneity, contributes to treatment efficacy in this tumor model.

### Simultaneous analysis of genetic and transcriptomic variation in single cell breast cancer

We next applied DENDRO to the analysis of data from a study of primary and metastasized breast cancer [20]. We focused on two tumors (BC03 and BC09) that had the most cells sequenced (Table S5 and Figure S12). BC03 included cells from the primary tumor (here after BC03P) as well as cells from regional metastatic lymph nodes (here after BC03LN), whereas BC09 included cells only from the primary resection. 132 single cell transcriptomes were profiled by Smart-seq protocol [17]. We first assess whether DENDRO separated BC03 and BC09, and then examine the transcriptomic and genetic heterogeneity within each tumor.

GATK [52] detected a total of 2,364,823 mutation sites across the 132 cells, 353,647 passed QC and were retained for downstream analysis (Figure S12A, B, C). Figure 4 shows the clustering determined by DENDRO. DENDRO separates BC09 cells from BC03 cells with 100% accuracy (Figure 4A). The intra-cluster divergence curve flattened at five subclones: three subclones for BC03 and two for BC09 (Figure 4A, Figure S12D and Table S3B). Within BC03, Clone Mix_1 and Clone Mix_2 contained a mixture of cells from the primary tumor and lymph nodes, and Clone LN_1 contained mostly cells from the lymph nodes. This suggests that tumor cells that have metastasized to the lymph nodes are genetically heterogeneous, with some cells remaining genetically similar to the primary population and others acquiring new genetic mutations. In comparison, hierarchical clustering based on expression (using log transcripts-per-million values) did not separate BC03 from BC09, and gave a negative adjusted Rand index within BC03, indicating effectively random assignment of cells to the two spatial locations (Figure 4B).

**Figure 4.**
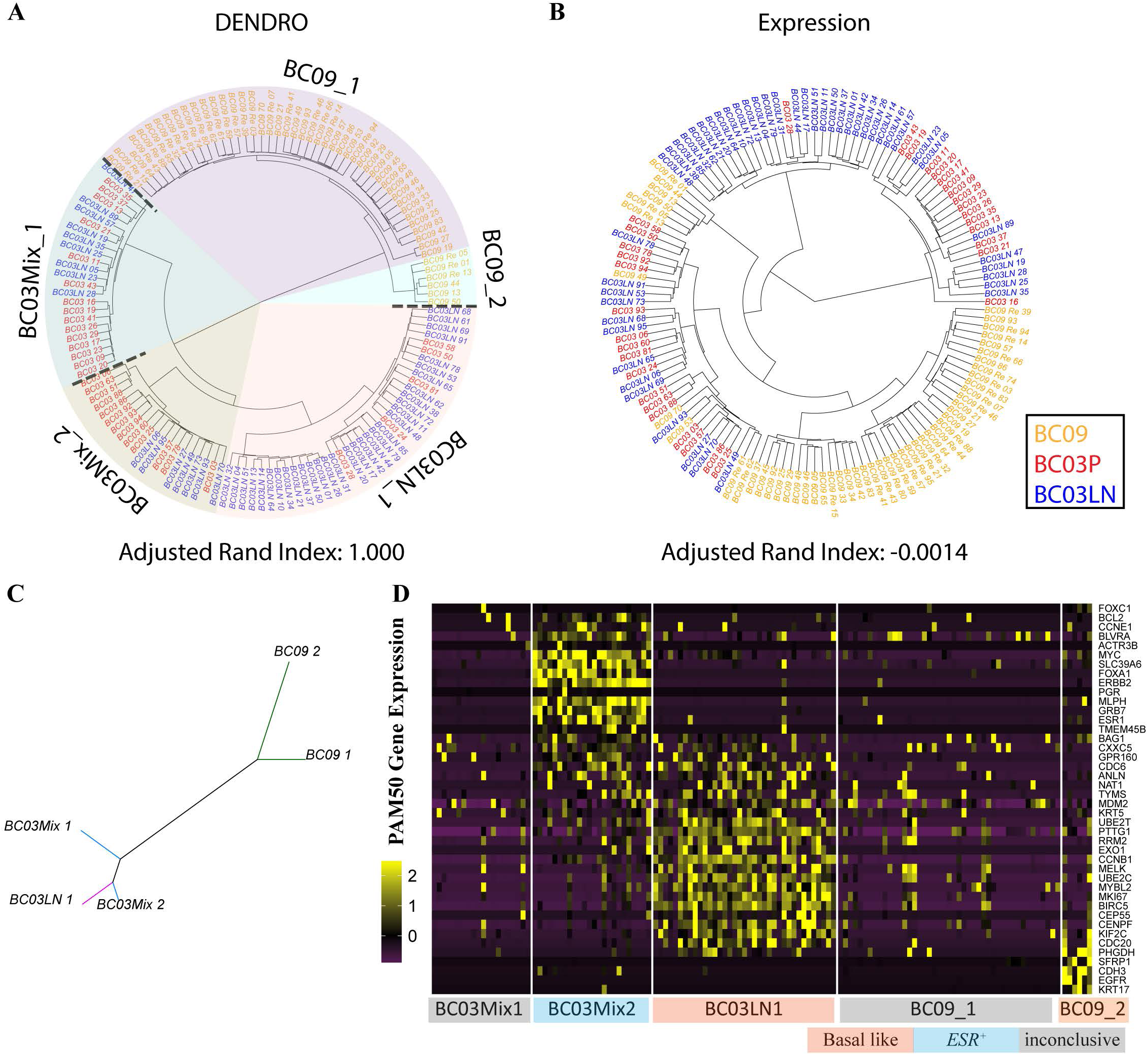
Analysis of scRNA-seq dataset of primary breast cancer. **a** DENDRO cluster result for primary breast cancer dataset (Chung et al., 2017). **b** Hierarchical clustering result for the same dataset based on expression (*log*TPM). (dashlines indicate cluster boundaries). **c** Neighbor joining phylogenetic tree given detected subclones for breast cancer dataset. **d** PAM50 gene panel expression shows breast cancer subtypes of each subclone.

We then pooled cells within each of the 5 clusters and re-estimated their mutation profiles with DENDRO. We defined a variant as subclonal if it was not present in all of the subclones within a tumor. Based on detection marginal likelihood, we picked the top 10,000 most confident variants to construct a phylogenetic tree (Figure 4C). As expected, the two BC09 clusters are far from the three BC03 clusters. Within BC03, the length of the branches shows that the subclone containing mostly cells from lymph nodes (labeled BC03LN_1) is genetically more similar to Clone Mix_2 compared to Clone Mix_1 (Figure 4C). In addition, window-smoothed expression plot with cells grouped by DENDRO clustering shows broad chromosome-level shifts in expression patterns between subclones, most likely due to copy number aberrations that are consistent with SNAs (Figure S13) [22].

A comparison of the transcriptomes of the subclones revealed substantial differences in the expression of PAM50 genes, which are prognostic markers for breast cancer (Figure 4D) [53]. DENDRO detected one rare subclone, BC09_2, with only six cells (<5% of the total number of cells) which had a strong basal-like signature. Interestingly, in BC03, Clone LN_1 has the TNBC/basal-like subtype with an invasive gene signature, while Clone Mix_2 has the *ESR1^+^* subtype. Thus, the genetic divergence of Clone LN_1 from Clone Mix_2 is accompanied by its acquisition of an invasive metastatic expression signature. In a direct comparison between cells from the primary site and cells from the lymph node without distinguishing subclones, these expression differences would be much weaker since the subclones do not cleanly separate by site. Compared with the original analysis that assigned each tumor to one specific breast cancer subtype, this analysis identifies subclones with different expression phenotypes, potentially allowing for better therapy design that targets all subclone phenotypes to reduce the risk of tumor relapse.

Existing scRNA-seq studies of cancer tissue cluster cells based on total gene expression or copy number profiles derived from smoothed total expression, making it difficult to separate the effects of sub-clonal copy number aberrations from transcriptomic variation [19, 22, 24]. Differential expression analysis based on clusters derived from total expression is prone to self-fulfilling prophecy, as there would indeed be differentially expressed genes because this is the clustering criteria. Because DENDRO’s subclone identification is based solely on genetic divergence, and not on expression profile, the downstream differential gene expression analysis can be precisely attributed to transcriptional divergence between subclones.

Hence, we conducted a transcriptome-wide search for pathways that have differential expression between subclones (Methods and Table S6), and assessed their overlap with pathways that are differentially mutated between subclones. Focusing on tumor BC03, pathways for G2M checkpoint and *KRAS* signaling are up-regulated in lymph node metastasis Clone BC03LN_1, while pathways for estrogen response and apoptosis are down-regulated, indicating a more invasive phenotype (Table S6E). In addition, *GAPDH* is up-regulated in the metastatic subclone (BC03LN_1) and down-regulated in the two mix-cell subclones, consistent with previous findings [54, 55] (Figure S14D). Differentially expressed genes between other subclone pairs in BC03 are also enriched in estrogen response, apoptosis, and DNA repair (Table S6C, D). In parallel, subclone-specific mutated genes are highly enriched in cancer-related pathways including MYC target, G2M checkpoints and mitotic spindle, and immune related pathways such as, interferon response, TNF-a signaling and inflammatory response (Table S6). Interestingly, few of the differentially mutated genes are associated with estrogen and androgen responses, suggesting that the differential expression of hormone related genes is not mediated directly by genetic mutations in these pathways. This is consistent with the recent studies that epigenetic alteration, such as histone acetylation and methylation, regulate hormones receptor signaling in breast cancer [56-59]. DNA-RNA joint analysis between other subclones are included in the Table S6 and Figure S14. Overall, this example illustrates how DENDRO enables the joint assessment of genetic and transcriptomic contributions to clonal diversity at single-cell resolution.

## Discussion

We have described DENDRO, a statistical framework to reconstruct intratumor DNA-level heterogeneity using scRNA-seq data. DENDRO starts with mutations detected directly from the scRNA-seq reads, which are very noisy due to a combination of factors: (1) errors are introduced in reverse-transcription, sequencing and mapping, (2) low sequencing depth and low molecule conversion efficiency leading to technical dropouts, and (3) expression burstiness at the single cell level leading to biological dropouts. DENDRO overcomes these obstacles through the statistical modeling of each component. Given noisy mutation profiles and allele-specific read counts, DENDRO computes a distance between each pair of cells that quantifies their genetic divergence after accounting for transcriptional bursting, dropout and sequencing error. Then, DENDRO clusters the cells based on this distance as subclone and re-estimates a more robust subclone-specific mutation profile by pooling reads across cells within the same cluster. These re-estimated mutations profiles are then passed to downstream mutation analysis and phylogenetic tree reconstruction.

Importantly, the genetic divergence used by DENDRO for cell clustering is based solely on allelic expression ratios and do not reflect the difference in total expression between cells at mutation sites. Thus, DENDRO differs from, and complements, existing tools that cluster cells based on total expression. In fact, as shown by spike-in analysis, DENDRO clusters the cells based on true underlining mutation profiles, and is robust to changes in total gene expression. As expected, the numbers of cells, the depth of sequencing, the actual number of subclonal mutations and the phylogenetic tree structure all influence the power of DENDRO. To aid researchers in experiment design, we developed DENDROplan, which predicts DENDRO’s clustering accuracy given basic experimental parameters and the expected informative mutation count, which can be obtained from bulk DNA sequencing.

Ideally, joint sequencing of the DNA and RNA on the same cells would allow us to relate genomic profiles to transcriptomic variations. Currently, there is yet no scalable technology for doing this. Separately performing scDNA-seq and scRNA-seq on different batches of cells within the same tumor would meet the nontrivial challenge of matching the subclones between the two data sets. DENDRO takes advantage of the central dogma and utilizes computational methods to extract genetic divergence information from noisy mutation calls in coding regions. Through two case studies, we illustrate the insights gained from the subclonal mutation and expression joint analysis that DENDRO enables.

We have demonstrated that proper computational modeling can excavate the DNA-level heterogeneity in scRNA-seq data. Yet, there is always limitation in working with RNA. While rare RNA editing events are absorbed by the parameter *ϵ*, DENDRO cannot distinguish subcluster-specific constituent RNA editing events from subcluster-specific DNA mutations. In the extreme and unlikely scenario where RNA editing events are common and pervasive, DENDRO’s cluster would reflect RNA editing. In such cases, we recommend using matched bulk DNA-seq of the same tumor to filter the loci detected in the first step of DENDRO, keeping only those that are supported by at least on read in the bulk DNA-seq data.

## Conclusions

We have developed DENDRO, a statistical method for tumor phylogeny inference and clonal classification using scRNA-seq data. DENDRO accurately infers the phylogeny relating the cells and assigns each single cell from the scRNA-seq data set to subclone. DENDRO allows us to (1) cluster cells based on genetic divergence while accounting for transcriptional bursting, technical dropout and sequencing error, as benchmarked by in silico mixture and a spike-in analysis, (2) characterize the transcribed mutations for each subclone, and (3) perform single-cell multi-omics analysis by examining the relationship between transcriptomic variation and mutation profile with the same set of cells. We evaluate the performance of DENDRO through a spike-in analysis and a data set with known subclonal structure. We further illustrate DENDRO through two case studies. In the first case study of relationship between intratumor heterogeneity and ICB treatment response, DENDRO estimates tumor mutation burden and predicts repertoire of high-affinity neoantigens in each subclone from scRNA-seq. In the second case study on a primary breast tumor dataset, DENDRO brought forth new insights on the interplay between intratumor transcriptomic variation and subclonal divergence.

## Methods

### scRNA-seq alignment and SNA calling pipeline

Figure S1 illustrates the SNA calling pipeline. Raw scRNA-seq data is aligned by STAR 2-pass method, which accounts for splicing junctions and achieve higher mapping quality [60]. In the next step, raw variants calling is made using the Haplotype Caller (GATK tool) on the BAM files after sorting, joining read groups, removing duplicated reads, removing overhangs into intronic regions, realigning and recalibration [61]. Conventionally, there are two methods from GATK tools for mutation detection: haplotype caller and mutect2. Haplotype caller has a RNA-seq setting which handle splice junctions correctly, but assumes VAF around 50%, while mutect2 can detect mutations with low VAF but does account for splice junction. The reason we select haplotype caller instead of mutect2 is that we extract allele read counts for all cells as long as one of the cells is listed as carrying the mutation. Thus, as long as one cell has VAF reaching 50%, this mutation would be detected. Calls with stand_call_conf greater than 20 and population frequency greater than 5% but less than 95% were preserved for further analysis. Admittedly, such lenient filtering typically introduces false positive sites. However, our priority at this step is to minimize false negative rate, while the genetic divergence matrix in the following step robustly estimates cell population substructure. Both the coverage of the alternative allele and the total read coverage are extracted for each site for further analysis.

### Genetic Divergence and Beta-Binomial framework

Consider two cells: *c* and *c*’. Let *I*_*c*_ and *I*_*c*′_ denote the clonal group to which the cells belong, i.e. *I*_*c*_ = *I*_*c*′_ if and only if cells *c* and *c*’ come from the same subclone. We define the genetic divergence at loci *g*, by 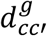:

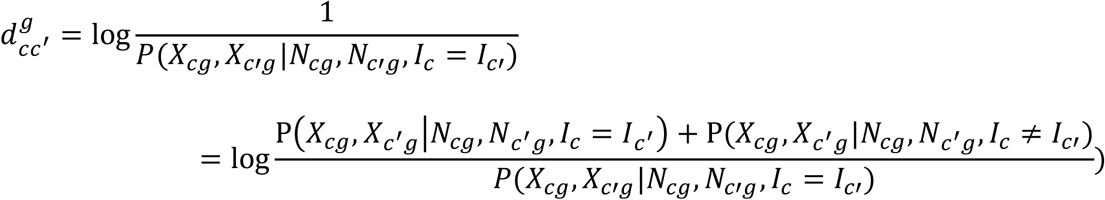

where *X*_*c*_ = (*X*_*c*1_, *X*_*c*2_,… *X*_*cg*_,…, *X*_*cm*_) are the mutation allele read counts for cell *c* and *N*_*c*_ = (*N*_*c*1_, *N*_*c*2_,… *N*_*cg*_,…, *N*_*cm*_) are the total read counts at these sites. More intuitively, if cells *c* and *c*′ are not from the same clonal group, the numerator has larger value compared to denominator. Thus 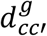 is large, indicating bigger divergence between the two cells. With further derivation (Supplementary Materials), 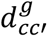 is a function of the five following probabilities:

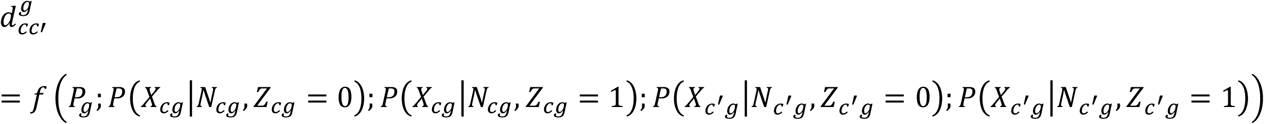

where *Z*_*cg*_ ∈ {0,1} is SNA indicator for cell *c* at site *g* and *P*_*g*_ = *P*(*Z*_*g*_ = 1) is mutation frequency across the cells estimated by GATK calls.

In the above formula for 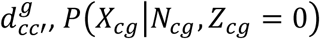 and *P*(*X*_*c*′*g*_|*N*_*cg*_,*Z*_*cg*_ = 0) reflect reverse-transcription/sequencing/mapping errors and rare RNA editing events, because when there is no mutation (i.e. *Z*_*cg*_ = 0), all mutation reads reflect such technical errors or RNA editing. Let ϵ denote the combined rate of technical error and RNA editing, we have

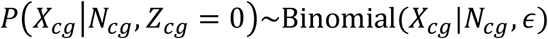

For cases where there are mutations (i.e. *Z*_*cg*_ = 1), the distribution of mutated read counts given total read counts is modeled with a Beta Binomial distribution, which is capable of modeling technical dropout and transcriptional bursting, and is supported by previous allele specific expression studies [30, 62].

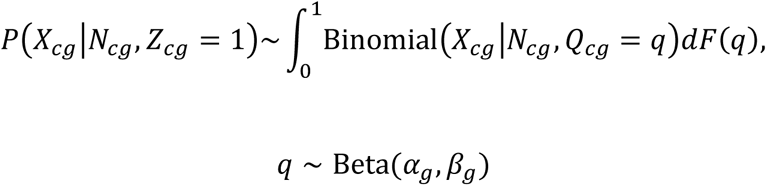

where *Q*_*cg*_ indicates proportion of mutated alleles expressed in cell *c* at site *g*, with Beta distribution as prior. Respectively, *α*_*g*_ and *β*_*g*_ represent gene activation and deactivation rate, which are estimated empirically.

### Kernel based clustering and optimal cluster assignment

We cluster the cells using a kernel-based algorithm, such as hierarchical clustering. Given that there are multiple sorting schemes, we leave the user to choose it. For the default-sorting scheme, we recommend “ward.D” [63]. This is because *d*_*cc*′_ behaves like a log likelihood ratio, which should follow a *χ*^2^ distribution when the two cells share the same subclone. The “ward.D” method has been shown to work well in Euclidian space. Empirically, among different hierarchical clustering algorithms on the renal cell carcinoma dataset (Figure S5) “ward.D” based hierarchical clustering performs the best.

To determine the number of clusters we use an intra-cluster divergence curve computed from the divergence matrix. Existing software rely on AIC, BIC, or another model selection metric [64, 65]. However, since we only have the “distance” matrix, these traditional methods cannot be applied. Let *N*_*k*_ be the number of cell pairs in cluster *C*_*k*_ and *N* be the total number of pairs between cells for all clusters. Let *K* be the number of clusters. The weighted sum of intra-cluster distance *W*_*K*_ is

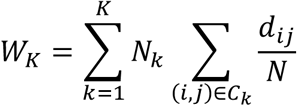

Note that small clusters are naturally down-weighted in the above metric. DENDRO relies on visual examination of the intra-cluster divergence curve (*W*_*K*_ plotted against *K*) to find the “elbow point”, which can be taken as a reasonable clustering resolution.

### Power analysis toolkit and experimental design

Before conducting a single cell RNA-seq experiment on a tumor sample, it is important to project how subclone detection power depends on the number of cells sequenced and the coverage per cell. To facilitate experiment design, we have developed a tool, DENDROplan (Figure 2a), that predicts the expected clustering accuracy by DENDRO given sequencing parameters and available bulk data for the tumor. Given an assumed tree structure and a target accuracy, DENDROplan computes the necessary read depth and number of cells needed.

As shown in Figure 2a, if bulk DNA sequencing and/or RNA sequencing data are available for the tumor being studied, these data can be harnessed to make more realistic power calculations. For example, if SNAs have been profiled using bulk DNA sequencing data, the set of mutations that lie in the exons of annotated genes can be retrieved and used directly in constructing the spike-in data. Furthermore, phylogeny construction algorithms for bulk DNA-seq data can be used to infer a putative tree structure that can be used as input to DENDROplan [5, 65]. If bulk RNA-seq data is available, the bulk expression level of the mutation-carrying genes can be used to predict the expression level of the mutation in the single cell data. For details, see Methods. The power analysis tool is also available at https://github.com/zhouzilu/DENDRO.

### Spike-in analysis

In our spike-in analysis, we adopt a dataset from Deng et al, which has allele specific read counts [39]. We further added binomial noise to mimic sequencing error. When analyzing DENDRO performance in terms of various number mutation site, number of cells, proportion of cells in each clade and proportion of mutations along each branch, we only take a subset of cells with similar cell types. On the other hand, we utilize a mixture cell population to test the robustness of DENDRO performance with regard to various expression profiles.

### SNA inference in “bulk” and phylogenetic tree construction

As stated previously, DENDRO further inferred SNA after pooling the reads from all cells within each cluster. Because, with our choice of thresholds, we identify SNAs in single cells with high sensitivity, the “bulk” level SNAs should be a subset of the SNAs in single cells, and mutation allele counts and total allele counts should provide us with enough information for SNA detection using a maximum likelihood framework [66], which accounts for both sequencing error and rare RNA-editing events. Suppose *g* is the genotype (number of reference allele) at a site and assume *m*, the ploidy, equals to 2. Then the likelihood is:

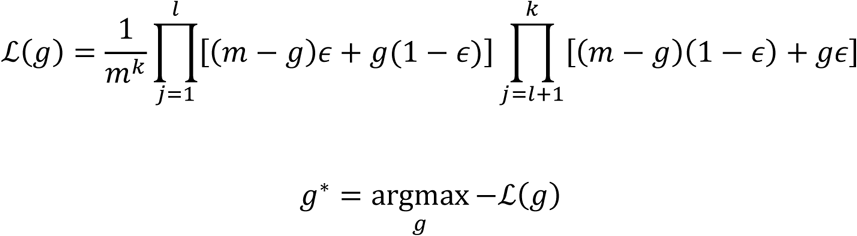

where *k* is number of reads at a site and the first *l* bases (*l* ≤ k) be the same to reference and the rests are same to alternative allele. E is the sequencing error and rare RNA-editing combined rate. Given mutation profiles, DENDRO then constructs a phylogenetic tree with the neighbor-joining method, which can more accurately capture the evolutionary relationship between different subclones [67] than the initial tree given by hierarchical clustering.

### Differential gene expression, mutation annotation and gene ontology analysis

We use Seurat and scDD to identify differentially expressed genes between tumors and between tumor subclones [68-70]. For each comparison, we apply two different methods: MAST implemented by Seurat and scDD. Genes with adjusted p-value < 0.05 count as significant differentially expressed gene for each method. We further intersect these two sets of differentially expressed genes to increase robustness. Subclonal mutations are annotated by ANNOVAR with default parameters and variants associated with intergenic regions were discarded for downstream analysis [71]. For GO analysis, we apply Gene Set Enrichment Analysis tool [52]. Hallmark gene sets serve as fundamental database with FDR q-value < 0.05 as significant.

### Single cell RNA-seq of Tumor Model Derived from B16

Six C57bl/6 mice were injected on both flanks with either B16 or R499: four with B16 and two with R499. Two of the mice implanted with B16 were treated with 200 ug of anti-PD1 per mouse on Days 5, 8 and 11. On Day 15, all tumors were harvested and made into single cell suspension. 100,000 CD45 negative tumor cells were sorted on Aria to enrich for live tumor cells and loaded on SMARTer ICELL8 cx Single-Cell System prior to full length single cell RNA-sequencing library preparation using Smart-seq following manufacturer’s recommendations. 460 cells and 11531 genes passed standard QC and were retained for downstream analysis.

### Neoantigen prediction

Based on gene expression from RNA-seq data, variants from unexpressed transcripts are removed. The MHC-I binding affinities of variants are then predicted using NetMHC version 4.0 for H-2-Kb and H-2-Db using peptide lengths from 8 to 11 [72]. Given subclonal mutation profile, we further assign the neoantigens to each subclone.

## List of abbreviations

SNA: Single nucleotide alteration
scDNA-seq: Single-cell DNA sequencing
scRNA-seq: Single-cell RNA sequencing
PDX: Patient-derived xenograft
TPM: Transcripts per kilobase million
ICB: Immune checkpoint blockade
TMB: Tumor mutational burden

## Declarations

### Acknowledgements

We thank Dr. Nan Lin for informing useful tools for biological insight analysis, and Dr. Kai Tan and Dr. Mingyao Li for helpful comments and suggestions.

### Availability of data and materials

DENDRO is an open-source R package available at https://github.com/zhouzilu/DENDRO with license GPL-3.0.

### Authors’ contributions

ZZ and NRZ formulated the model. ZZ developed and implemented the algorithm and conducted all computational analyses. BX and AM designed the immunotherapy case study. BX performed the experiments of the immunotherapy case. ZZ, BX and NRZ wrote the manuscript. All authors read and approved the final manuscript.

### Consent for publication

Not applicable.

### Ethics approval and consent to participate

Not applicable.

## References

1. Gamazon, E.R. and B.E. Stranger, The impact of human copy number variation on gene expression. Briefings in Functional Genomics, 2015. 14(5): p. 352-357.

2. Hanks, S., et al., Constitutional aneuploidy and cancer predisposition caused by biallelic mutations in BUB1B. Nat Genet, 2004. 36(11): p. 1159–61.

3. Vicente-Duenas, C., et al., Epigenetic Priming in Cancer Initiation. Trends Cancer, 2018. 552 4(6): p. 408–417.

4. Burrell, R.A., et al., The causes and consequences of genetic heterogeneity in cancer evolution. Nature, 2013. 501(7467): p. 338–45.

5. Jiang, Y., et al., Assessing intratumor heterogeneity and tracking longitudinal and spatial clonal evolutionary history by next-generation sequencing. Proc Natl Acad Sci U S A, 557 2016. 113(37): p. E5528–37.

6. Deshwar, A.G., et al., PhyloWGS: reconstructing subclonal composition and evolution from whole-genome sequencing of tumors. Genome Biol, 2015. 16: p. 35.

7. Zare, H., et al., Inferring clonal composition from multiple sections of a breast cancer. PLoS Comput Biol, 2014. 10(7): p. e1003703.

8. Carter, S.L., et al., Absolute quantification of somatic DNA alterations in human cancer. Nat Biotechnol, 2012. 30(5): p. 413-21.

9. Li, B. and J.Z. Li, A general framework for analyzing tumor subclonality using SNP array and DNA sequencing data. Genome Biol, 2014. 15(9): p. 473.

10. Oesper, L., A. Mahmoody, and B.J. Raphael, THetA: inferring intra-tumor heterogeneity from high-throughput DNA sequencing data. Genome Biol, 2013. 14(7): p. R80.

11. Ha, G., et al., TITAN: inference of copy number architectures in clonal cell populations from tumor whole-genome sequence data. Genome Res, 2014. 24(11): p. 1881–93.

12. Miller, C.A., et al., SciClone: inferring clonal architecture and tracking the spatial and temporal patterns of tumor evolution. PLoS Comput Biol, 2014. 10(8): p. e1003665.

13. Navin, N.E., The first five years of single-cell cancer genomics and beyond. Genome Res, 573 2015. 25(10): p. 1499–507.

14. Navin, N., et al., Tumour evolution inferred by single-cell sequencing. Nature, 2011. 575 472(7341): p. 90–4.

15. Wang, Y., et al., Clonal evolution in breast cancer revealed by single nucleus genome sequencing. Nature, 2014. 512(7513): p. 155–60.

16. Gao, R., et al., Punctuated copy number evolution and clonal stasis in triple-negative breast cancer. Nat Genet, 2016. 48(10): p. 1119–30.

17. Picelli, S., et al., Smart-seq2 for sensitive full-length transcriptome profiling in single cells. Nat Methods, 2013. 10(11): p. 1096-8.

18. Klein, A.M., et al., Droplet barcoding for single-cell transcriptomics applied to embryonic stem cells. Cell, 2015. 161(5): p. 1187–1201.

19. Patel, A.P., et al., Single-cell RNA-seq highlights intratumoral heterogeneity in primary glioblastoma. Science, 2014. 344(6190): p. 1396–401.

20. Chung, W., et al., Single-cell RNA-seq enables comprehensive tumour and immune cell profiling in primary breast cancer. Nat Commun, 2017. 8: p. 15081.

21. Kim, K.T., et al., Application of single-cell RNA sequencing in optimizing a combinatorial therapeutic strategy in metastatic renal cell carcinoma. Genome Biol, 2016. 17: p. 80.

22. Tirosh, I., et al., Dissecting the multicellular ecosystem of metastatic melanoma by single-cell RNA-seq. Science, 2016. 352(6282): p. 189–96.

23. Jerby-Arnon, L., et al., A Cancer Cell Program Promotes T Cell Exclusion and Resistance to Checkpoint Blockade. Cell, 2018. 175(4): p. 984–997 e24.

24. Tirosh, I., et al., Single-cell RNA-seq supports a developmental hierarchy in human oligodendroglioma. Nature, 2016. 539(7628): p. 309–313.

25. Venteicher, A.S., et al., Decoupling genetics, lineages, and microenvironment in IDH-mutant gliomas by single-cell RNA-seq. Science, 2017. 355(6332).

26. Li, H., et al., Reference component analysis of single-cell transcriptomes elucidates cellular heterogeneity in human colorectal tumors. Nat Genet, 2017. 49(5): p. 708-718.

27. van Galen, P., et al., Single-Cell RNA-Seq Reveals AML Hierarchies Relevant to Disease 601 Progression and Immunity. Cell, 2019. 176(6): p. 1265–1281 e24.

28. Nam, A.S., et al., Somatic mutations and cell identity linked by Genotyping of Transcriptomes. Nature, 2019. 571(7765): p. 355–360.

29. Raj, A., et al., Stochastic mRNA synthesis in mammalian cells. PLoS Biol, 2006. 4(10): p. 605 e309.

30. Jiang, Y., N.R. Zhang, and M. Li, SCALE: modeling allele-specific gene expression by single-cell RNA sequencing. Genome Biol, 2017. 18(1): p. 74.

31. Padovan-Merhar, O., et al., Single mammalian cells compensate for differences in 609 cellular volume and DNA copy number through independent global transcriptional 610 mechanisms. Mol Cell, 2015. 58(2): p. 339–52.

32. Zafar, H., et al., Monovar: single-nucleotide variant detection in single cells. Nat 612 Methods, 2016. 13(6): p. 505–7.

33. Piskol, R., G. Ramaswami, and J.B. Li, Reliable identification of genomic variants from RNA-seq data. Am J Hum Genet, 2013. 93(4): p. 641–51.

34. Brennecke, P., et al., Accounting for technical noise in single-cell RNA-seq experiments. Nat Methods, 2013. 10(11): p. 1093-5.

35. Pierson, E. and C. Yau, ZIFA: Dimensionality reduction for zero-inflated single-cell gene expression analysis. Genome Biol, 2015. 16: p. 241.

36. Vallejos, C.A., J.C. Marioni, and S. Richardson, BASiCS: Bayesian Analysis of Single-Cell Sequencing Data. PLoS Comput Biol, 2015. 11(6): p. e1004333.

37. Ding, B., et al., Normalization and noise reduction for single cell RNA-seq experiments. Bioinformatics, 2015. 31(13): p. 2225-7.

38. Qiu, X., et al., Single-cell mRNA quantification and differential analysis with Census. Nat 624 Methods, 2017. 14(3): p. 309–315.

39. Deng, Q., et al., Single-cell RNA-seq reveals dynamic, random monoallelic gene expression in mammalian cells. Science, 2014. 343(6167): p. 193–6.

40. Eirew, P., et al., Dynamics of genomic clones in breast cancer patient xenografts at single-cell resolution. Nature, 2015. 518(7539): p. 422–6.

41. Gerlinger, M., et al., Intratumor heterogeneity and branched evolution revealed by multiregion sequencing. N Engl J Med, 2012. 366(10): p. 883–892.

42. Shi, Y.J., et al., Intratumoral Heterogeneity in Breast Cancer: A Comparison of Primary and Metastatic Breast Cancers. Oncologist, 2017. 22(4): p. 487–490.

43. Ribas, A. and J.D. Wolchok, Cancer immunotherapy using checkpoint blockade. Science, 634 2018. 359(6382): p. 1350–1355.

44. Schumacher, T.N. and R.D. Schreiber, Neoantigens in cancer immunotherapy. Science, 636 2015. 348(6230): p. 69–74.

45. Rizvi, N.A., et al., Cancer immunology. Mutational landscape determines sensitivity to PD-1 blockade in non-small cell lung cancer. Science, 2015. 348(6230): p. 124–8.

46. Tumeh, P.C., et al., PD-1 blockade induces responses by inhibiting adaptive immune 640 resistance. Nature, 2014. 515(7528): p. 568–71.

47. Twyman-Saint Victor, C., et al., Radiation and dual checkpoint blockade activate non-redundant immune mechanisms in cancer. Nature, 2015. 520(7547): p. 373–7.

48. Benci, J.L., et al., Opposing Functions of Interferon Coordinate Adaptive and Innate Immune Responses to Cancer Immune Checkpoint Blockade. Cell, 2019. 178(4): p. 933-645 948 e14.

49. Patel, S.A. and A.J. Minn, Combination Cancer Therapy with Immune Checkpoint Blockade: Mechanisms and Strategies. Immunity, 2018. 48(3): p. 417–433.

50. Goodman, A.M., et al., Tumor Mutational Burden as an Independent Predictor of Response to Immunotherapy in Diverse Cancers. Mol Cancer Ther, 2017. 16(11): p. 2598–650 2608.

51. Rosenthal, R., et al., Neoantigen-directed immune escape in lung cancer evolution. Nature, 2019. 567(7749): p. 479-485.

52. Subramanian, A., et al., Gene set enrichment analysis: a knowledge-based approach for 654 interpreting genome-wide expression profiles. Proc Natl Acad Sci U S A, 2005. 102(43): p. 655 15545–50.

53. Parker, J.S., et al., Supervised risk predictor of breast cancer based on intrinsic subtypes. J 657 Clin Oncol, 2009. 27(8): p. 1160–7.

54. Zhang, J.Y., et al., Critical protein GAPDH and its regulatory mechanisms in cancer cells. Cancer Biol Med, 2015. 12(1): p. 10-22.

55. Tarrado-Castellarnau, M., et al., Glyceraldehyde-3-phosphate dehydrogenase is overexpressed in colorectal cancer onset. Translational Medicine Communications, 2017. 662 2(1): p. 6.

56. Mann, M., V. Cortez, and R.K. Vadlamudi, Epigenetics of estrogen receptor signaling: role in hormonal cancer progression and therapy. Cancers (Basel), 2011. 3(3): p. 1691-707.

57. Green, K.A. and J.S. Carroll, Oestrogen-receptor-mediated transcription and the influence 666 of co-factors and chromatin state. Nat Rev Cancer, 2007. 7(9): p. 713–22.

58. Dreijerink, K.M., et al., Menin links estrogen receptor activation to histone H3K4 trimethylation. Cancer Res, 2006. 66(9): p. 4929–35.

59. Kim, H., et al., Requirement of histone methyltransferase SMYD3 for estrogen receptor-mediated transcription. J Biol Chem, 2009. 284(30): p. 19867–77.

60. Dobin, A., et al., STAR: ultrafast universal RNA-seq aligner. Bioinformatics, 2013. 29(1): 672 p. 15–21.

61. McKenna, A., et al., The Genome Analysis Toolkit: a MapReduce framework for analysing next-generation DNA sequencing data. Genome Res, 2010. 20(9): p. 1297–303.

62. Skelly, D.A., et al., A powerful and flexible statistical framework for testing hypotheses of allele-specific gene expression from RNA-seq data. Genome Res, 2011. 21(10): p. 1728-677 37.

63. Ward, J.H., Hierarchical Grouping to Optimize an Objective Function. Journal of the 679 American Statistical Association, 1963. 58(301): p. 236-&.

64. Goutte, C., et al., Feature-space clustering for fMRI meta-analysis. Hum Brain Mapp, 681 2001. 13(3): p. 165–83.

65. Urrutia, E., et al., Integrative pipeline for profiling DNA copy number and inferring tumor phylogeny. Bioinformatics, 2018. 34(12): p. 2126–2128.

66. Li, B., et al., A likelihood-based framework for variant calling and de novo mutation detection in families. PLoS Genet, 2012. 8(10): p. e1002944.

67. Schliep, K.P., phangorn: phylogenetic analysis in R. Bioinformatics, 2011. 27(4): p. 592-3.

68. Finak, G., et al., MAST: a flexible statistical framework for assessing transcriptional changes and characterizing heterogeneity in single-cell RNA sequencing data. Genome 689 Biol, 2015. 16: p. 278.

69. Korthauer, K.D., et al., A statistical approach for identifying differential distributions in single-cell RNA-seq experiments. Genome Biol, 2016. 17(1): p. 222.

70. Butler, A., et al., Integrating single-cell transcriptomic data across different conditions, technologies, and species. Nat Biotechnol, 2018. 36(5): p. 411–420.

71. Wang, K., M. Li, and H. Hakonarson, ANNOVAR: functional annotation of genetic 695 variants from high-throughput sequencing data. Nucleic Acids Res, 2010. 38(16): p. 696 e164.

72. Karosiene, E., et al., NetMHCcons: a consensus method for the major histocompatibility complex class I predictions. Immunogenetics, 2012. 64(3): p. 177–86.

