## Supplementary Material for "Genetic Heterogeneity Profiling by Single Cell RNA Sequencing"

*Derivation of Beta-Binomial framework*

Consider two cells: $c$ and $c'$. Let $I_c$ and $I_{c\prime}$ denote the clonal group of the cells, i.e. $I_c = I_{c\prime}$ indicates cell $c$ and $c'$ come from same subclone. Genetic divergence $d_{cc\prime}$ is the negative log likelihood of the mutation allele counts of cells $c$ and $c'$, given the total read counts and the event that the two cells belong to the same subclone.

$$d_{cc'} = d_{cc\prime} = \log P(X_c, X_{c'} | N_c, N_{c'}, I_c = I_{c'}) = \sum_{g=1}^{m} d_{cc\prime}^{g}$$

Given $d_{cc'}^{g} = -\log P\left(X_{cg}, X_{c'g} \middle| N_{cg}, N_{c'g}, I_c = I_{c'}\right)$,

$$d_{cc'}^{g} = \log \frac{1}{P\left(X_{cg}, X_{c'g} \middle| N_{cg}, N_{c'g}, I_c = I_{c'}\right)}$$

$$= \log \frac{\mathrm{P}\left(X_{cg}, X_{c'g} \middle| N_{cg}, N_{c'g}, I_c = I_{c'}\right) + \mathrm{P}\left(X_{cg}, X_{c'g} \middle| N_{cg}, N_{c'g}, I_c \neq I_{c'}\right)}{P\left(X_{cg}, X_{c'g} \middle| N_{cg}, N_{c'g}, I_c = I_{c'}\right)})$$

$$= \log(\frac{P(X_{cg}, X_{cg'} | N_{cg}, N_{cg'}, I_c \neq I_{c\prime})}{P(X_{cg}, X_{cg'} | N_{cg}, N_{cg'}, I_c = I_{c\prime})} + 1)$$

where $D_c = \{(N_{c1}, N_{c2}, \dots N_{cg}, \dots N_{cm}), (X_{c1}, X_{c2}, \dots X_{cg}, \dots X_{cm})$ are data for cell $c$.

$\frac{P(X_{cg}, X_{cg'} | N_{cg}, N_{cg'}, I_c \neq I_{c\prime})}{P(X_{cg}, X_{cg'} | N_{cg}, N_{cg'}, I_c = I_{c\prime})}$ could also be called a Bayes Factor. More intuitively, if cell $c$ and $c'$ are not from the same clonal group, the numerator has larger value compared to denominator. Thus, $d_{cc\prime}^{g}$ is large, indicating bigger divergence between the two cells.

To further expand the formula, let us focus on the denominator first:

$$P\left(X_{cg}, X_{c'g} \,|N_{cg}, N_{c'g}, I_c = I_{c'}\right)$$

$$= P\left(X_{cg}, X_{c'g} \,|N_{cg}, N_{c'g}, Z_{cg} = Z_{c'g} = 0\,\right)P\left(Z_{cg} = Z_{c'g} = 0|I_{cg} = I_{c'g}\right)$$
$$+ P\left(X_{cg}, X_{c'g} \,|N_{cg}, N_{c'g}, Z_{cg} = Z_{c'g} = 1\,\right)P\left(Z_{cg} = Z_{c'g} = 1|I_{cg} = I_{c'g}\right)$$

$$= P\left(X_{cg} \,|N_{cg},\, Z_{cg} = 0\right)P\left(X_{c'g} \,|N_{c'g},\, Z_{c'g} = 0\right)\left(1 - P_g\right)$$
$$+ P\left(X_{cg} \,|N_{cg},\, Z_{cg} = 1\right)P\left(X_{c'g} \,|N_{c'g},\, Z_{c'g} = 1\right)P_g$$

Then the numerator:

$$P\left(X_{cg}, X_{c'g} \,|N_{cg}, N_{c'g}, I_c \neq I_{c'}\right)$$

$$= P\left(X_{cg}, X_{c'g} \,|N_{cg}, N_{c'g}, Z_{cg} = 0,\, Z_{c'g} = 0\,\right)P\left(Z_{cg} = 0, Z_{c'g} = 0|I_{cg} \neq I_{c'g}\right)$$
$$+ P\left(X_{cg}, X_{c'g} \,|N_{cg}, N_{c'g}, Z_{cg} = 1,\, Z_{c'g} = 1\,\right)P\left(Z_{cg} = 1, Z_{c'g} = 1|I_{cg} \neq I_{c'g}\right)$$
$$+ P\left(X_{cg}, X_{c'g} \,|N_{cg}, N_{c'g}, Z_{cg} = 0,\, Z_{c'g} = 1\,\right)P\left(Z_{cg} = 0, Z_{c'g} = 1|I_{cg} \neq I_{c'g}\right)$$
$$+ P\left(X_{cg}, X_{c'g} \,|N_{cg}, N_{c'g}, Z_{cg} = 1,\, Z_{c'g} = 0\,\right)P\left(Z_{cg} = 1, Z_{c'g} = 0|I_{cg} \neq I_{c'g}\right)$$

$$= P\left(X_{cg} \,|N_{cg},\, Z_{cg} = 1\right)P\left(X_{c'g} \,|N_{c'g},\, Z_{c'g} = 0\right)\left(1 - P_g\right)P_g$$
$$+ P\left(X_{cg} \,|N_{cg},\, Z_{cg} = 0\right)P\left(X_{c'g} \,|N_{c'g},\, Z_{c'g} = 0\right)\left(1 - P_g\right)^2$$
$$+ P\left(X_{cg} \,|N_{cg},\, Z_{cg} = 1\right)P\left(X_{c'g} \,|N_{c'g},\, Z_{c'g} = 1\right)P_g^2$$
$$+ P\left(X_{cg} \,|N_{cg},\, Z_{cg} = 0\right)P\left(X_{c'g} \,|N_{c'g},\, Z_{c'g} = 1\right)\left(1 - P_g\right)P_g$$

Thus, we have $d_{cc'}^{g}$ is a function of the following five probabilities

$$d_{cc'}^{g} = f\left(P_g; P\left(X_{cg}|N_{cg}, Z_{cg} = 0\right); P\left(X_{cg}|N_{cg}, Z_{cg} = 1\right); P\left(X_{c'g}|N_{cg}, Z_{cg} = 0\right); P\left(X_{c'g}|N_{cg}, Z_{cg} = 1\right)\right)$$

*Quantitative function analysis on genetic divergence evaluation by simulation*

Better understanding of the genetic divergence evaluation function is essential to DENDRO. Especially, we need to make sure DENDRO is capturing DNA level information from mutation rather than RNA level information from relative expression. This function, however, is quite complicated and difficult to analyze directly. As a result, we design several simulation schemes and analyze the function performance given different variables.

Let's consider 2 cells: cell $c$ and $c'$. We design true mutation profile $Z$ (an indicator function, 1 means mutation and 0 means no mutation) and relative read count $\theta$ (relative number by $\frac{\theta_i}{\Sigma\theta_j}$) at each position as Figure S2.

We further define true mutation distance $d_z$ as

$$d_z = \frac{|Z_c - Z_{c'}|_1}{m} = p$$

where $m$ is the total number of mutations. In another word, $d_z = p$, which is the true mutation rate in cell $c$.

We also define $\Delta = |H - M| = |M - L|$, representing relative expression differences.

Then, the relative expression distance $d_\theta$ can be written as,

$$d_\theta = \sum_i \frac{|\theta_i - \theta_{i'}|_1}{m} = (1-p)\Delta$$

We further have genetic divergence, $d_L$, calculated by negative log likelihood in DENDRO.

It is also interested in studying the relationship with $N$, the total number of read counts for each cell.

In our simulation, we alter $\Delta, p$ and $N$, assessing the responses in $d_L, d_\theta$ and $d_z$. Results show that (1) our genetic divergence function is orthogonal to relative expression level (Figure S2B). With fixed mutation rate $p$ and total read counts $N$, as relative expression differences $d_\theta$ increases, $d_L$ stay constant; (2) When we keep relative expression differences $d_\theta$ and total read counts $N$ as constant, as number of true mutation ($d_z$ or $p$) increases, genetic divergence $d_L$ also increases (Figure S2C); and (3) genetic divergence $d_L$ monotonically increases with total expression level ($N$) if we fix mutation distance $d_z$ (Figure S2D). This can be interpreted that with more reads support, DENDRO is more confident that the two cells belong to different clusters.

*DENDRO spike-in analysis*

A systematic evaluation of DENDRO's subclone detection accuracy on Deng et al. shows: (1) As mutation burden of a clade increases, the adjusted Rand index and clade-specific capture rate decrease, but cluster purity of interested clade increases (Figure S3A), because there are few mutations remaining to distinguish the other two subclones; (2) similarly, as cell proportion of a clade increases, its adjusted Rand index and capture rate decrease, but cluster purity increases (Figure S3B); (3) more mutation site increases DENDRO performance (Figure S3); (4) lower sequencing depth decreases accuracy (Figure S3); (5) As false positive rate of mutation calls increasing, DENDRO performs worse; (6) lower sequencing error rate improves DENDRO accuracy; and (7) variation in total expression of genes does not influence DENDRO's divergence measure (Figure S3C).

*DENDROplan evaluation metrics*

We evaluate DENDRO accuracy in DENDROplan with three different metrics: Adjusted Rand index, capture rate and purity.

1. Adjusted Rand index: Adjusted Rand index is a measure of the similarity between two data clusterings after adjusted for the chance grouping of elements. For details, see https://en.wikipedia.org/wiki/Rand_index
2. Capture rate: Capture rate is a measure of "false negative rate" of a specific clade. Out of all the cells from the specific clade, how many of them is detected by the algorithm.
3. Purity: Purity is a measure of "false positive rate" of a specific clade. Out of all the cells in the "specific cluster" you detected, how many are actually from the true specific clade.

*DENDRO analysis of a renal cell carcinoma and its metastasis*

GATK detected 2,867,029 mutation sites across all cells [1]. Mutations that are detected in less than 5% (too rare) or more than 95% (too common) of the cells were removed, which leaves 72,206 mutations. On average, 10801 mutations are detected in each cell and 17.35 cells possess the same mutation for each loci (Figure S4B, C). For majority sites, only few cells have nonzero read coverage, highlighting the fact that many mutations are missed due to technical and biological dropout (Figure S4D) [2-6].

After the cells were clustered into three subclones, DENDRO reesimate the clonal mutation profiles. By maximum likelihood, we called a total of 13,454 mutations after pooling the cells within each cluster (Figure S4F and Methods). The unrooted phylogenetic tree computed with the revised mutation profiles confirms that genetic divergence between metastasis and primary is greater than divergence between PDX and primary sample.

DENDRO enables simultaneous clonal assignment and transcriptomic profiling of the same set of cells. Plot of smoothed expression ordered by DENDRO shows unique expression patterns within each subclone (Figure S6). We focused on the comparison of the two metastasized cell populations (metastasis to lung and patient derived mouse xenograph). Even though PDX_mRCC was derived from Pt_mRCC, the DENDRO analysis found substantial genetic divergence between the two cell populations. To investigate further, we performed a differential expression analysis between PDX_mRCC and Pt_mRCC with scDD and MAST, detecting 74 significant differentially expressed genes (Methods, Figure S7E, Table S4A) [44-46]. Gene ontology analysis classified these 74 genes into two subgroups: immune-related genes and cancer-related genes (Table S1, Table S4F) [47]. Immune-related differentially expressed genes are enriched for the terms TNF-$\alpha$ signaling, complement system and allograft rejection. On the other hand, cancer related differentially expressed genes overlap with the pathways including hypoxia, KRAS signaling, mTORC1 signaling and epithelial mesenchymal transition.

Simultaneously, we compare the mutation profiles of these two subclones. 9521 locus have different mutated allele counts between these two populations and were further annotated by ANNOVAR [48]. After filtering, the preserved variants associated with 24 out of 74 differential expressed genes (Table S4A). Next, we performed a similar GSEA on variants associated genes to identify mutation-related pathway [47]. Interestingly, variant annotated genes are enriched in cancer-related pathways, including mitotic spindle, mTORC1 signaling, EMT and hypoxia, overlapping substantially with the cancer-related pathways identified by differential expression analysis; in comparison, none of the differentially expressed genes from immune pathways showed up in this mutated gene analysis (Table S2, Table S4F). In another word, cancer-related transcriptomic divergence between PDX_mRCC and Pt_mRCC is driven directly by genetic alterations in the same genes, but immune-related differential expression is influenced by non-DNA factors. This makes sense, since implantation of tumor cells from human to mice alters their immune microenvironment [49-51], and thus is expected to alter immune-related signaling within the implanted tumor cells. This illustrates how DENDRO extricates DNA variation from RNAs allowing heir joint analysis. Differential expression and differential mutation analysis for the other subclone pairs can be found in Table S4 and Figure S7.

Table S1. GO analysis on Differential Expressed Genes between Pt_mRCC and PDX_mRCC

| Gene Set Name [# Genes (K)] | | | | p-value | FDR q-value |
|---|---|---|---|---|---|
| **Color key:** | Cancer-related pathway | Immune-related pathway | Other | | |
| HALLMARK_TNFA_SIGNALING_VIA_NFKB [200] | | | | 1.86 e-8 | 9.28 e-7 |
| HALLMARK_HYPOXIA [200]* | | | | 5.04 e-7 | 1.26 e-5 |
| HALLMARK_MTORC1_SIGNALING [200]* | | | | 1.15 e-5 | 1.92 e-4 |
| HALLMARK_COMPLEMENT [200] | | | | 2.15 e-4 | 1.79 e-3 |
| HALLMARK_GLYCOLYSIS [200]* | | | | 2.15 e-4 | 1.79 e-3 |
| HALLMARK_KRAS_SIGNALING_UP [200] | | | | 2.15 e-4 | 1.79 e-3 |
| HALLMARK_ALLOGRAFT_REJECTION [200] | | | | 3.17 e-3 | 1.58 e-2 |
| HALLMARK_EPITHELIAL_MESENCHYMAL_TRANSINSITION [200]* | | | | 3.17 e-3 | 1.58 e-2 |
| HALLMARK_ESTROGEN_RESPONSE_EARLY [200] | | | | 3.17 e-3 | 1.58 e-2 |
| HALLMARK_INFLAMMATORY_RESPONSE [200] | | | | 3.17 e-3 | 1.58 e-2 |

Table S2. GO analysis on Differential Mutated Genes between Pt_mRCC and PDX_mRCC

| Gene Set Name [# Genes (K)] | | | | p-value | FDR q-value |
|---|---|---|---|---|---|
| **Color key:** | Cancer-related pathway | Immune-related pathway | Other | | |
| HALLMARK_UV_RESPONSE_DN [144] | | | | 1.39 e-29 | 6.93 e-28 |
| HALLMARK_MYC_TARGETS_V1 [200] | | | | 2.75 e-27 | 6.87 e-26 |
| HALLMARK_MITOTIC_SPINDLE [200] | | | | 7.97 e-24 | 1.33 e-22 |
| HALLMARK_MTORC1_SIGNALING [200]* | | | | 9.49 e-20 | 1.19 e-18 |
| HALLMARK_EPITHELIAL_MESENCHYMAL_TRANSINSITION [200]* | | | | 1.91 e-17 | 1.91 e-16 |
| HALLMARK_OXIDATIVE_PHOSPHORYLATION [200] | | | | 1.06 e-16 | 8.81 e-16 |
| HALLMARK_HYPOXIA [200]* | | | | 5.69 e-16 | 4.06 e-15 |
| HALLMARK_GLYCOLYSIS [200]* | | | | 2.97 e-15 | 1.86 e-14 |
| HALLMARK_ANDROGEN_RESPONSE [101] | | | | 1.07 e-14 | 5.96 e-14 |
| HALLMARK_HEME_METABOLISM [200] | | | | 7.4 e-14 | 3.7 e-13 |

Reference


1. McKenna, A., et al., *The Genome Analysis Toolkit: a MapReduce framework for analyzing next-generation DNA sequencing data.* Genome Res, 2010. **20**(9): p. 1297-303.
2. Brennecke, P., et al., *Accounting for technical noise in single-cell RNA-seq experiments.* Nat Methods, 2013. **10**(11): p. 1093-5.
3. Ding, B., et al., *Normalization and noise reduction for single cell RNA-seq experiments.* Bioinformatics, 2015. **31**(13): p. 2225-7.
4. Pierson, E. and C. Yau, *ZIFA: Dimensionality reduction for zero-inflated single-cell gene expression analysis.* Genome Biol, 2015. **16**: p. 241.
5. Vallejos, C.A., J.C. Marioni, and S. Richardson, *BASiCS: Bayesian Analysis of Single-Cell Sequencing Data.* PLoS Comput Biol, 2015. **11**(6): p. e1004333.
6. Qiu, X., et al., *Single-cell mRNA quantification and differential analysis with Census.* Nat Methods, 2017. **14**(3): p. 309-315.
