## Supplementary Figure for "Genetic Heterogeneity Profiling by Single Cell RNA Sequencing"

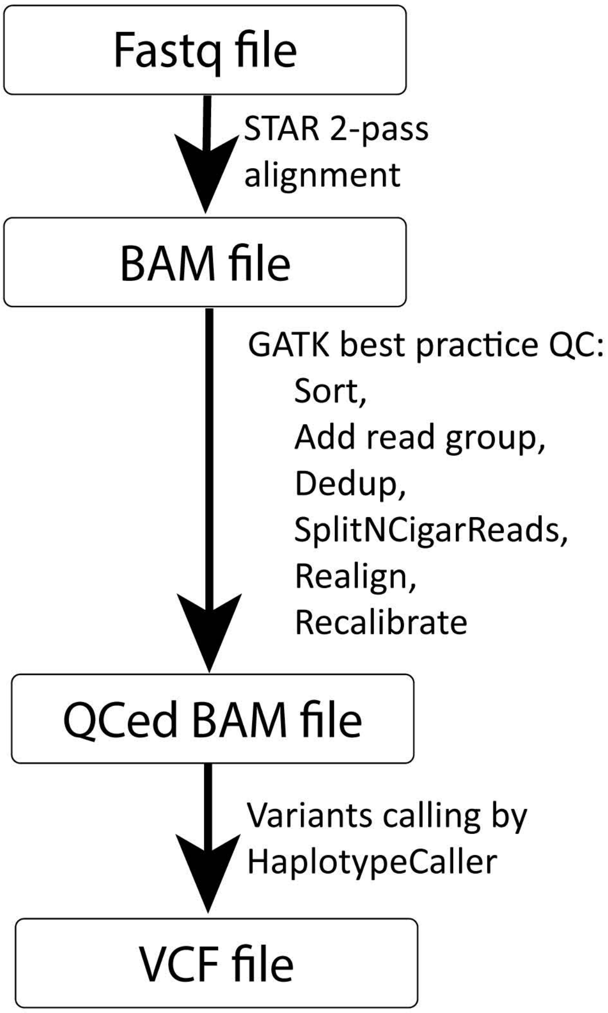


**Figure S1. An illustration of the SNA calling pipeline.** Raw scRNA-seq data is aligned by STAR 2-pass. Further quality control and variants calling steps follow GATK tool best practice by Broad Institute.

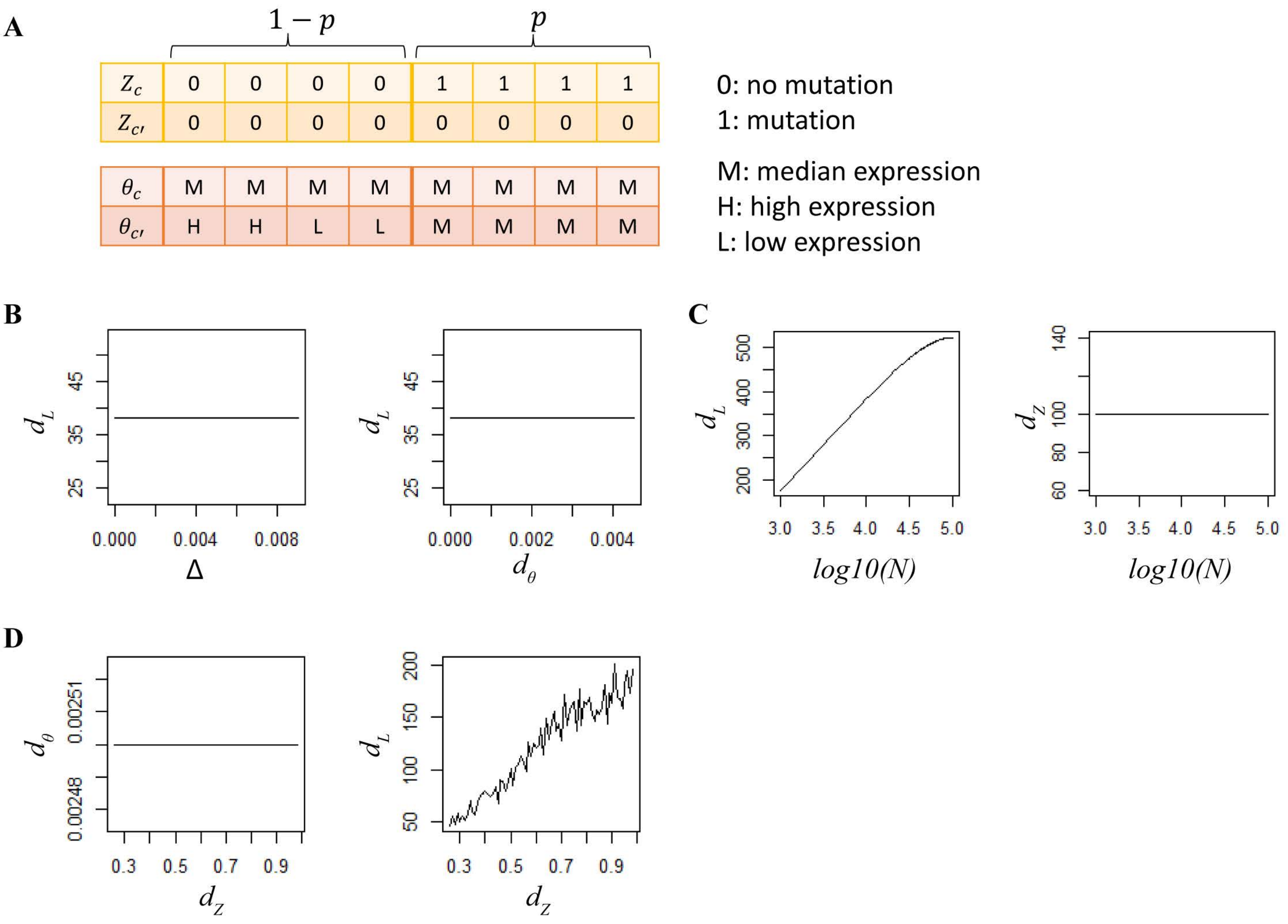


**Figure S2. Kernel function justification by simulation. a** Illustration of simulation set up. **b** With fixed *p* and *N*, as $d_\theta$ increases, $d_L$ stay constant. Thus, likelihood kernel is orthogonal to relative expression. **c** With fixed *p=1*, as *N* approach infinite, $d_L$ increase monotonically. As there are higher expression, we are more confident that the two cells belong to different clusters. **d** When $d_\theta$ and *N* stay the same, as $d_Z$ approaches 1, $d_L$ increases, because true mutation number increases.

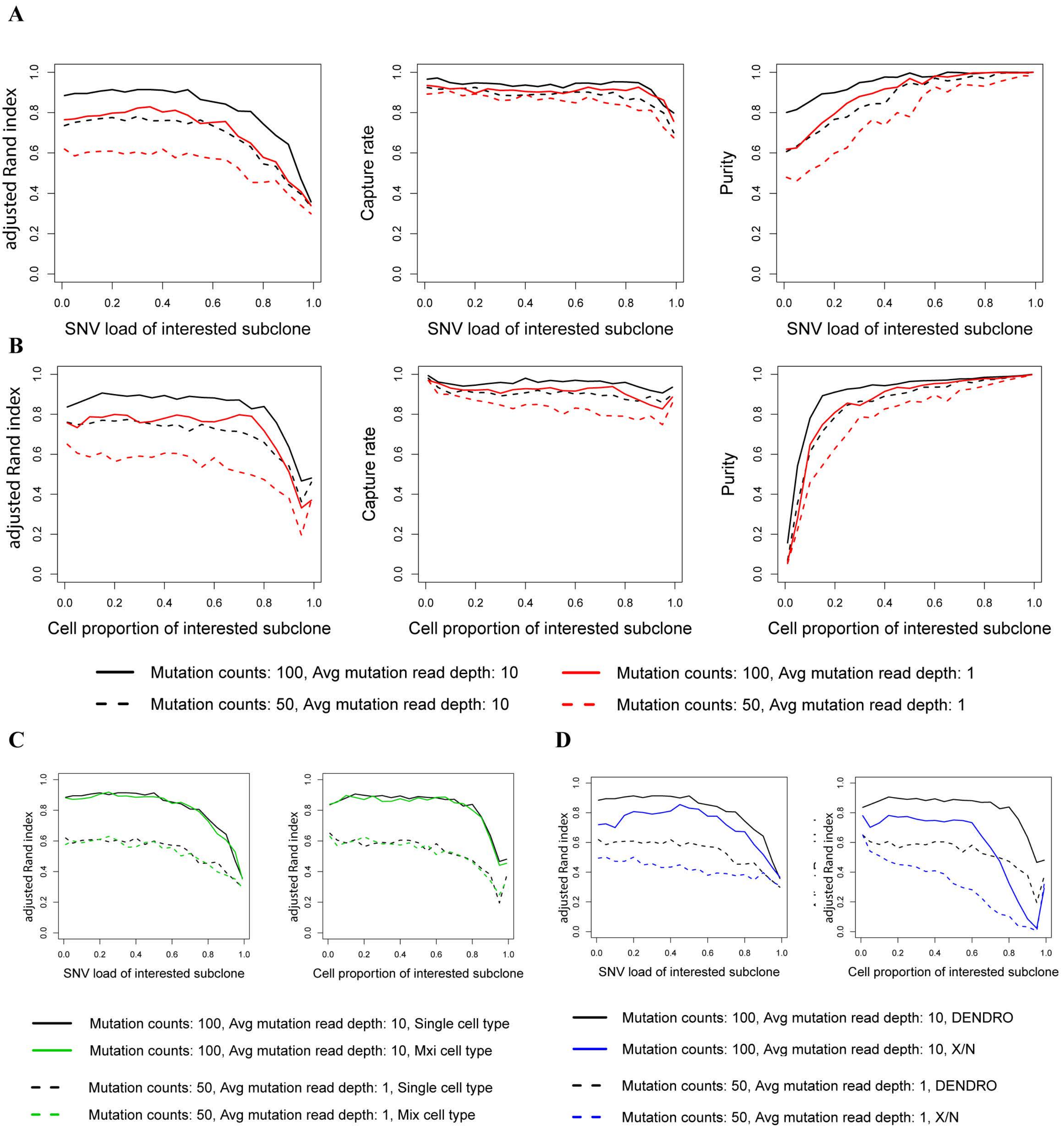


**Figure S3. DENDRO accuracy assessment by spike-in analysis. a** Statistics under different SNV load of interested clade (i.e. as fraction of total mutation counts) in a pure cell population vs. a mixture of two cell types (50% each). **b** Statistics under different cell proportion of interested clade in a pure cell population vs. a mixture of two cell types (50% each). (Mutation counts: number of mutation identified; Avg mutation read depth: average read depth of all the mutation sites.) Both plots show that mixture of cell population does not effect accuracy. **c** DENDRO accuracy assessment in mixture cell types. **d** Clustering accuracy assessment using DENDRO vs hclust on variants allele frequencies

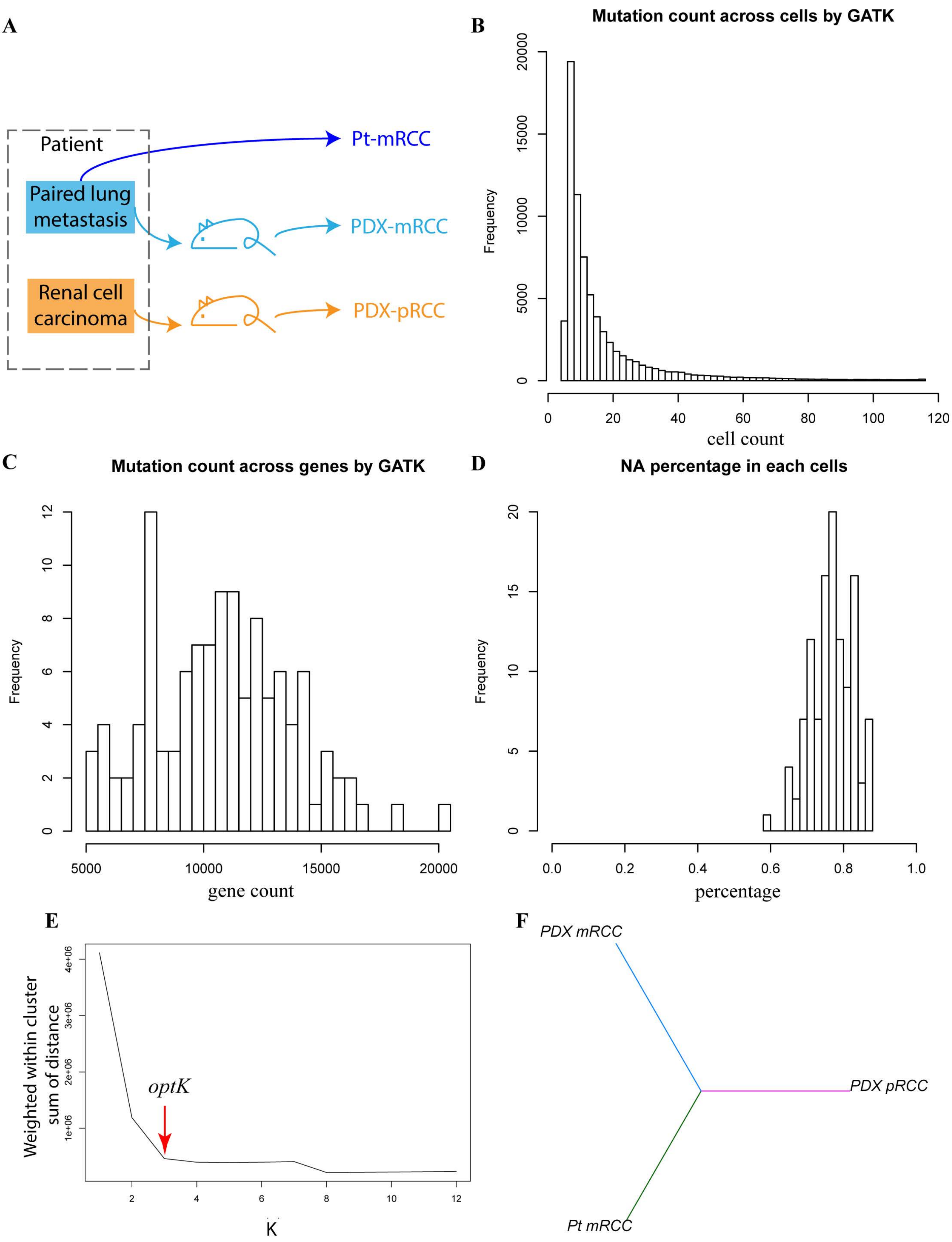


**Figure S4. RCC experimental design and its mutation statistics detected by GATK tool. a** Experimental design for renal cell carcinoma dataset. Figure modified from Kim et al. **b** Mutation count across cells by GATK. Most of the genes have low mutation frequency. **c** Mutation count across genes by GATK tool. It shows mutation counts with a bell shape. **d** NA percentage in each cell across genes. When there is no read counts, it shows as NA. **e** Intra-cluster divergence curve to select optimal number of cluster by DENDRO. Here, *optK=3* **f** Phylogenetic tree of three cell populations identified by DEDNRO. (In **a** - **d**, dashlines indicate cluster boundaries)

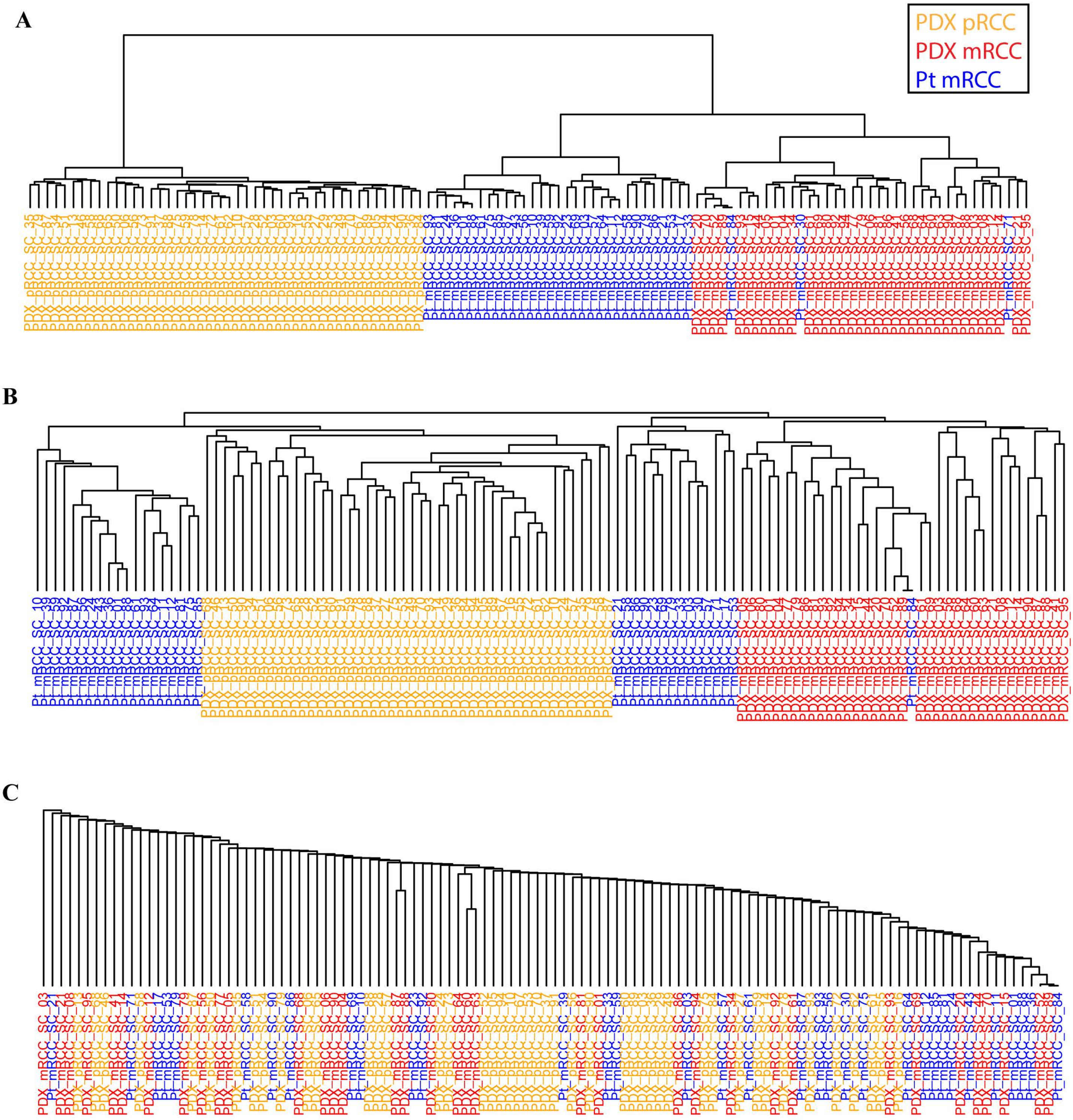


**Figure S5. Hierarchical clustering algorithm for renal cell carcinoma dataset. a** Genetic divergence matrix clustering by Ward.D algorithm. **b** Genetic divergence matrix clustering by Complete algorithm. **c** Genetic divergence matrix clustering by Single algorithm

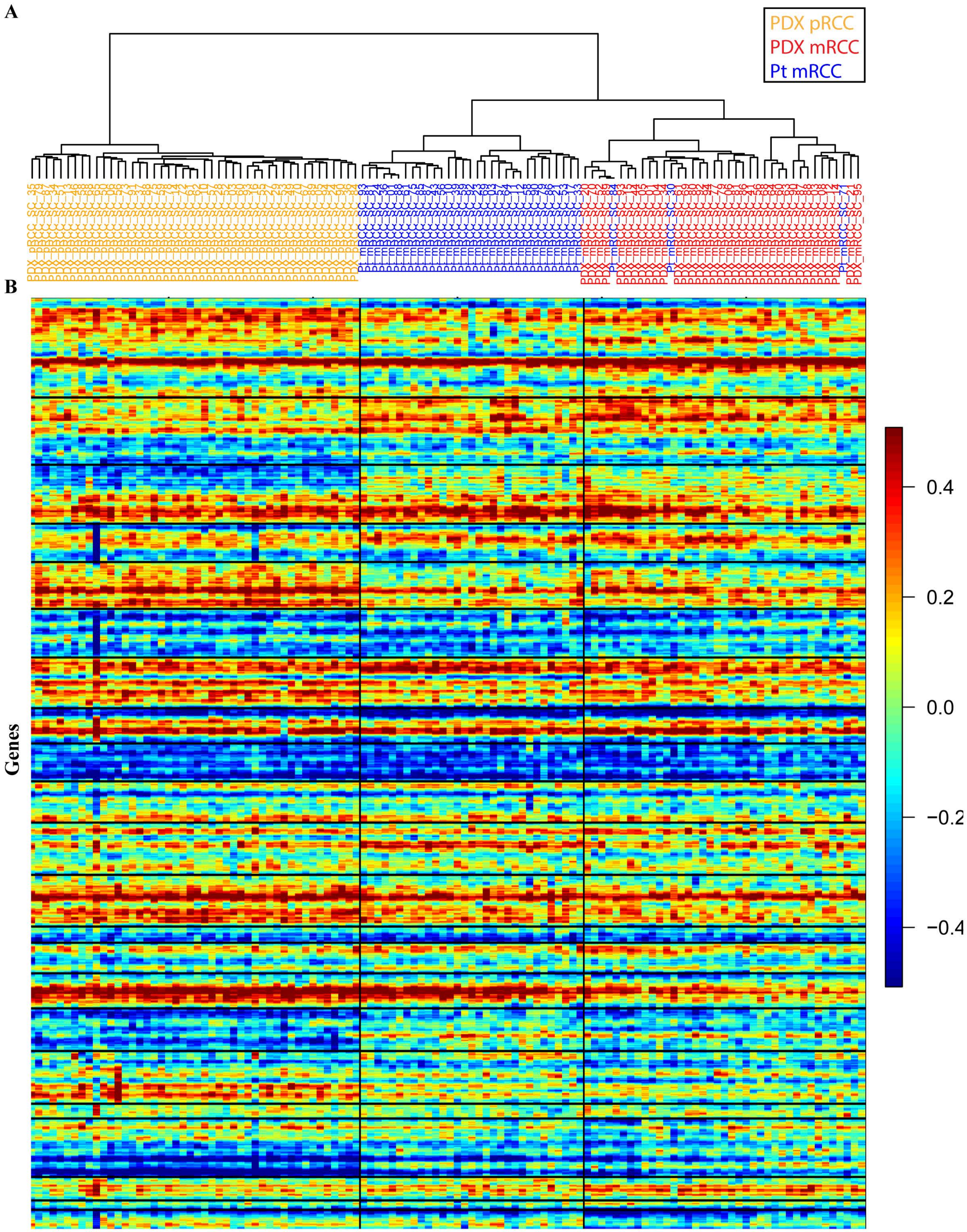


**Figure S6. Expression of renal cell carcinoma. a** DENDRO clustering of RCC. **b** Expression ordered by DENDRO clustering. Vertical line separate cluster identified by DEDNRO. Horizontal line separate different chromosomes.

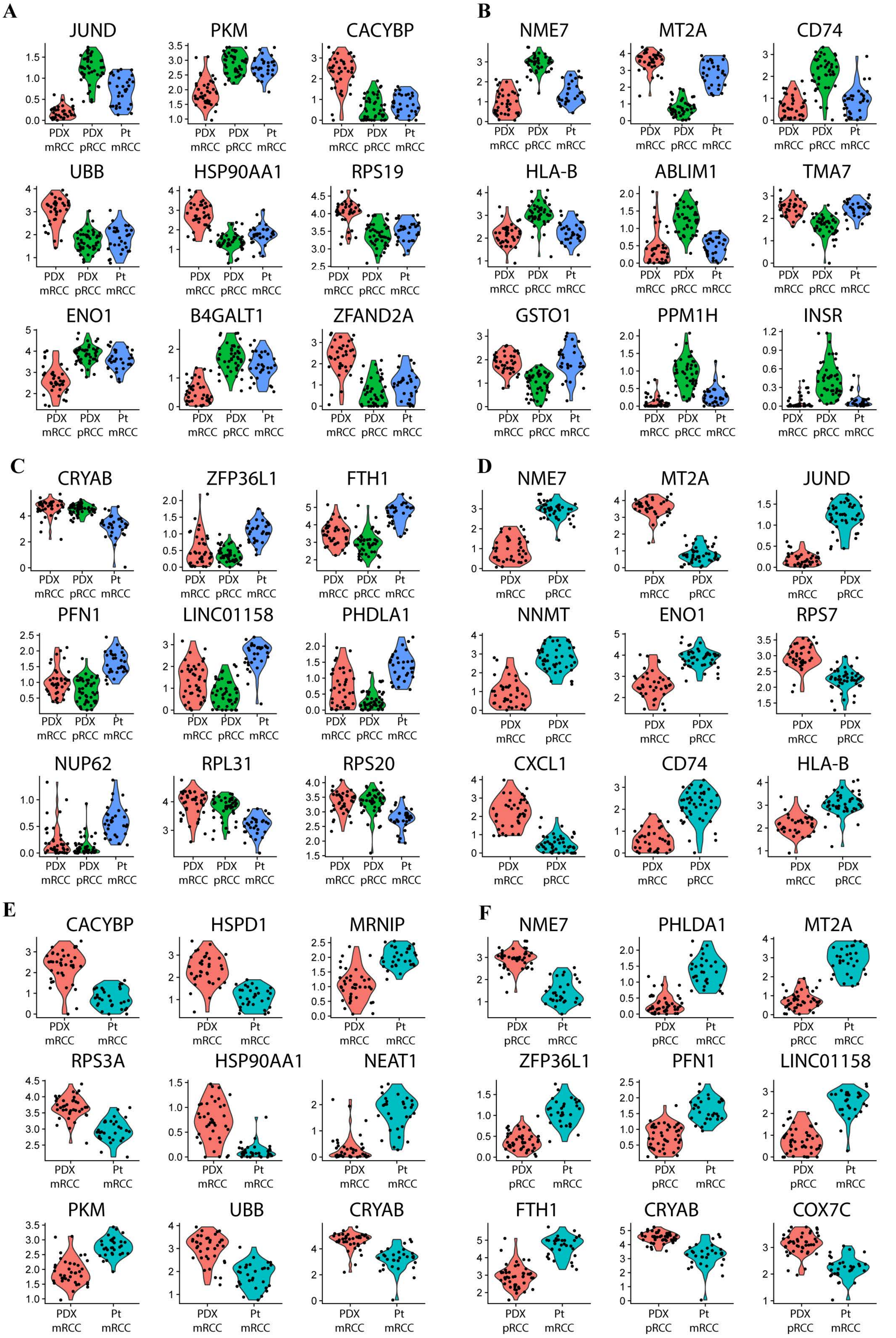


**Figure S7. Most significant differeitnal expressed genes between RCC pairs. a** PDX_mRCC vs. other. **b** PDX_pRCC vs. other. **c** Pt_mRCC vs. other. **d** PDX_mRCC vs. PDX_pRCC. **e** PDX_mRCC vs. Pt_mRCC. **f** Pt_mRCC vs. PDX_pRCC.

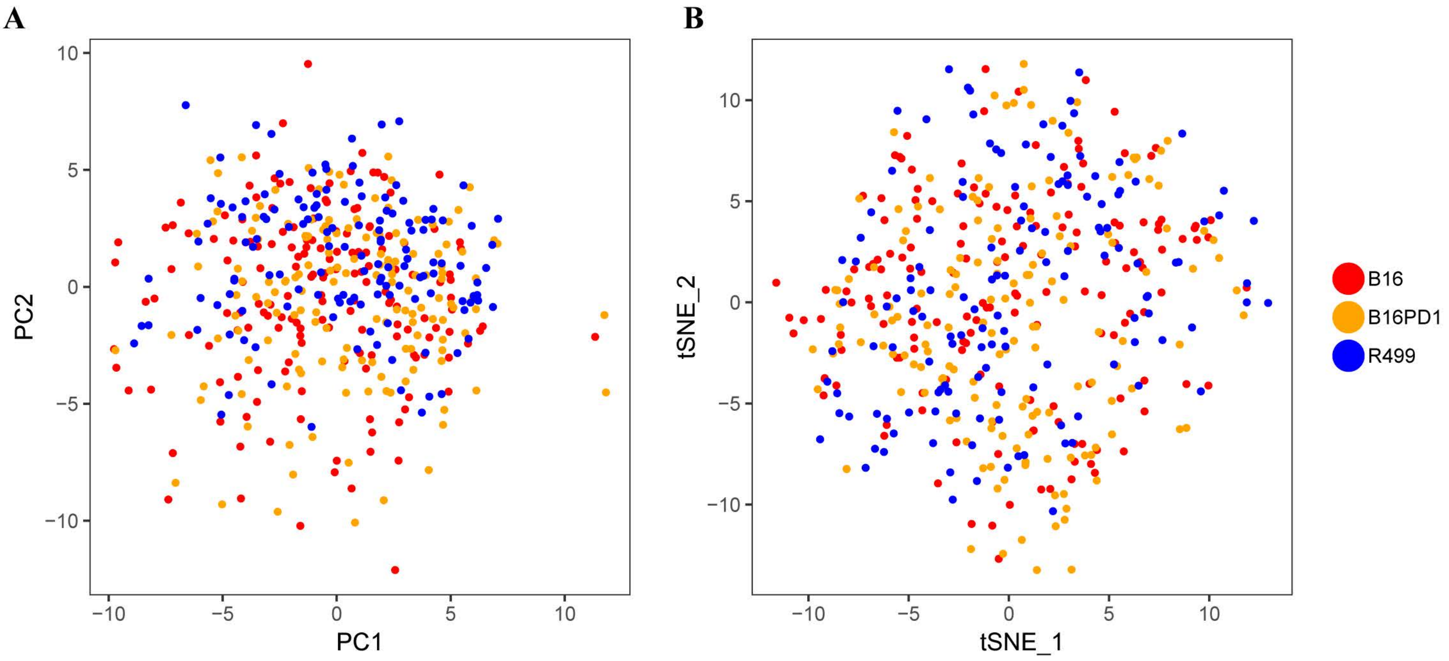


**Figure S8. Transcriptome analysis on anti-PD1 treatment experiment. a** PCA plot of the cells based on expression. **b** t-SNE plot of the cells based on expression.

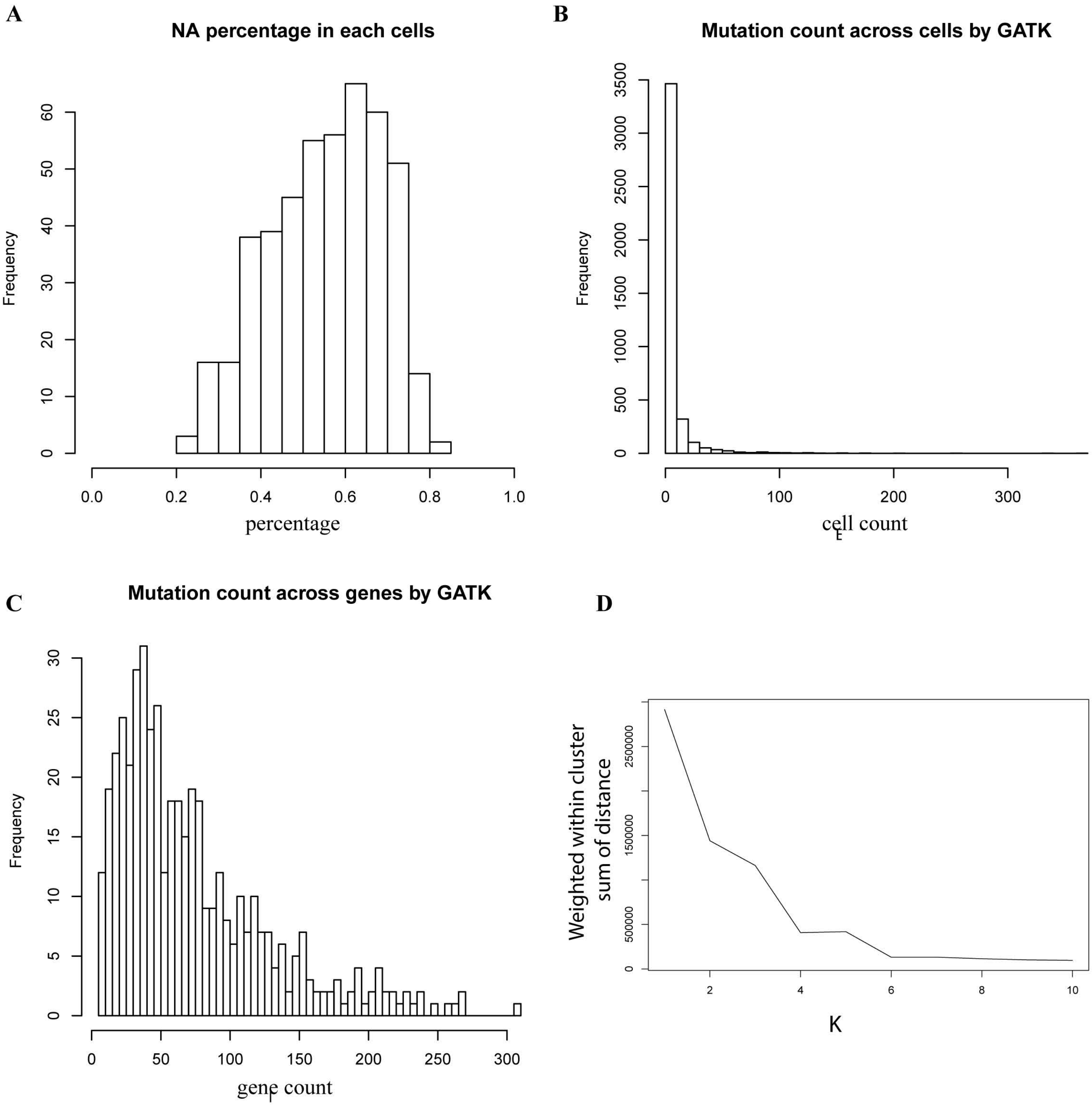


**Figure S9. Anti-PD1 treatment experiment mutation statistics detected by GATK tool and optimal clustering option. a** NA percentage in each cell across genes. When there is no read counts, it shows as NA. **b** Mutation count across cells by GATK. Most of the genes have low mutation frequency. **c** Mutation count across genes by GATK tool. It shows mutation counts with a bell shape. **d** Intra-cluster divergence curve given different number of clusters (K). 4 is the elbow point.

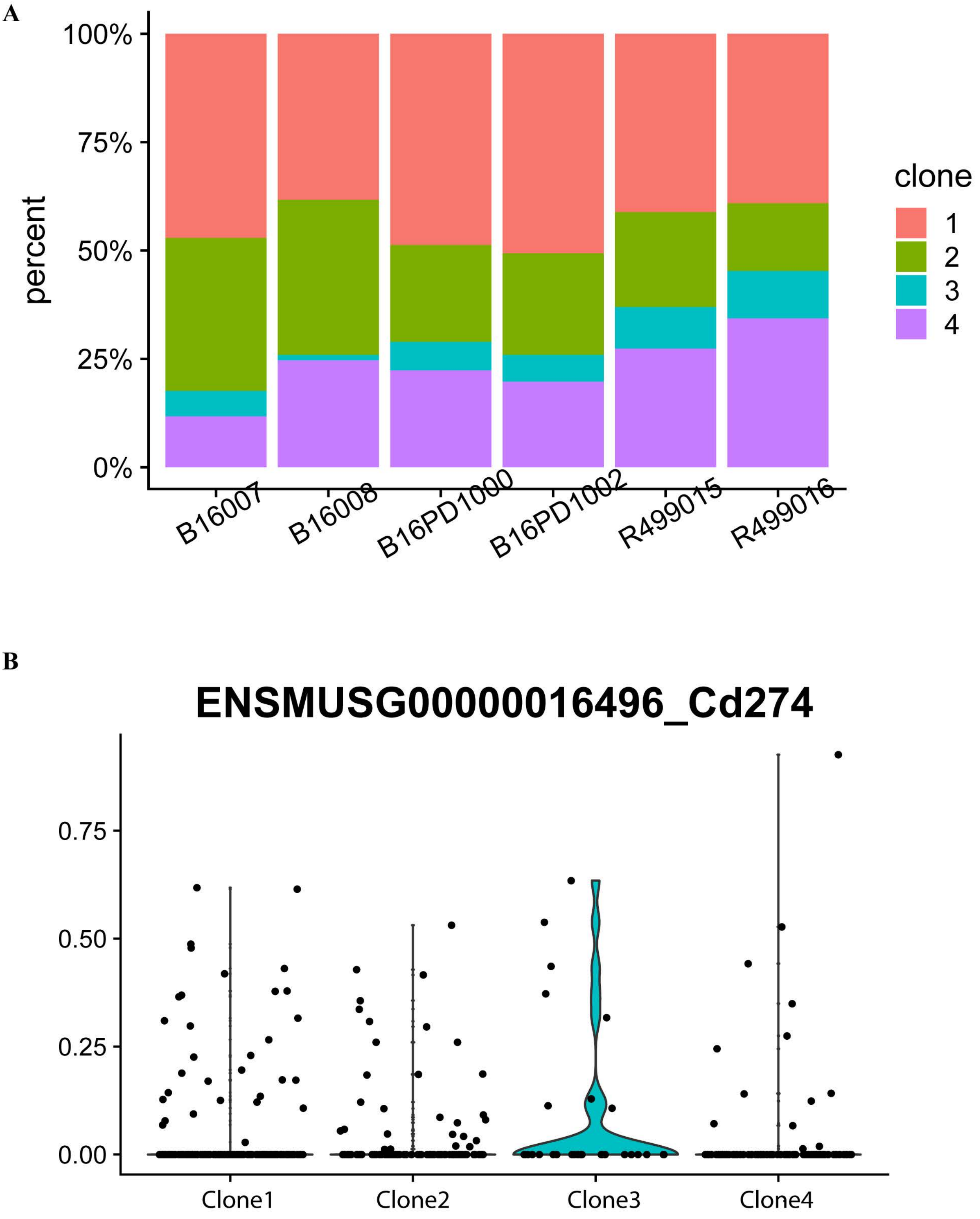


**Figure S10. Anti-PD1 treatment experiment. a** Frequencies of the subclonal population in each of the 6 tumor samples. **b** Expression level of *Pdl1* (*Cd274*) in each clone.

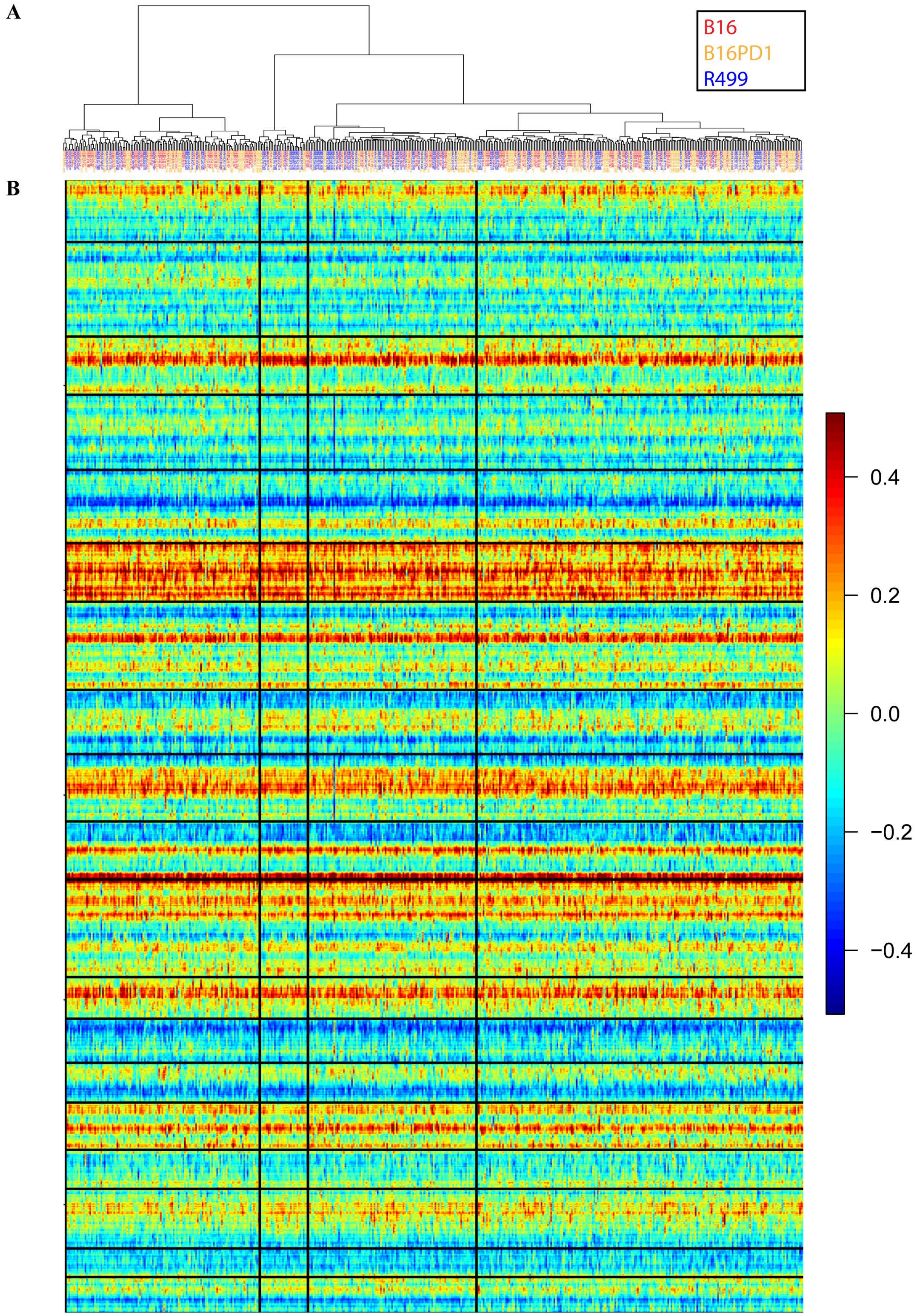


**Figure S11. Smoothed expression of anti-PD1 treatment experiment a** DENDRO clustering. Color indicates various conditions. **b** Smoothed expression heatmap ordered by DENDRO clustering

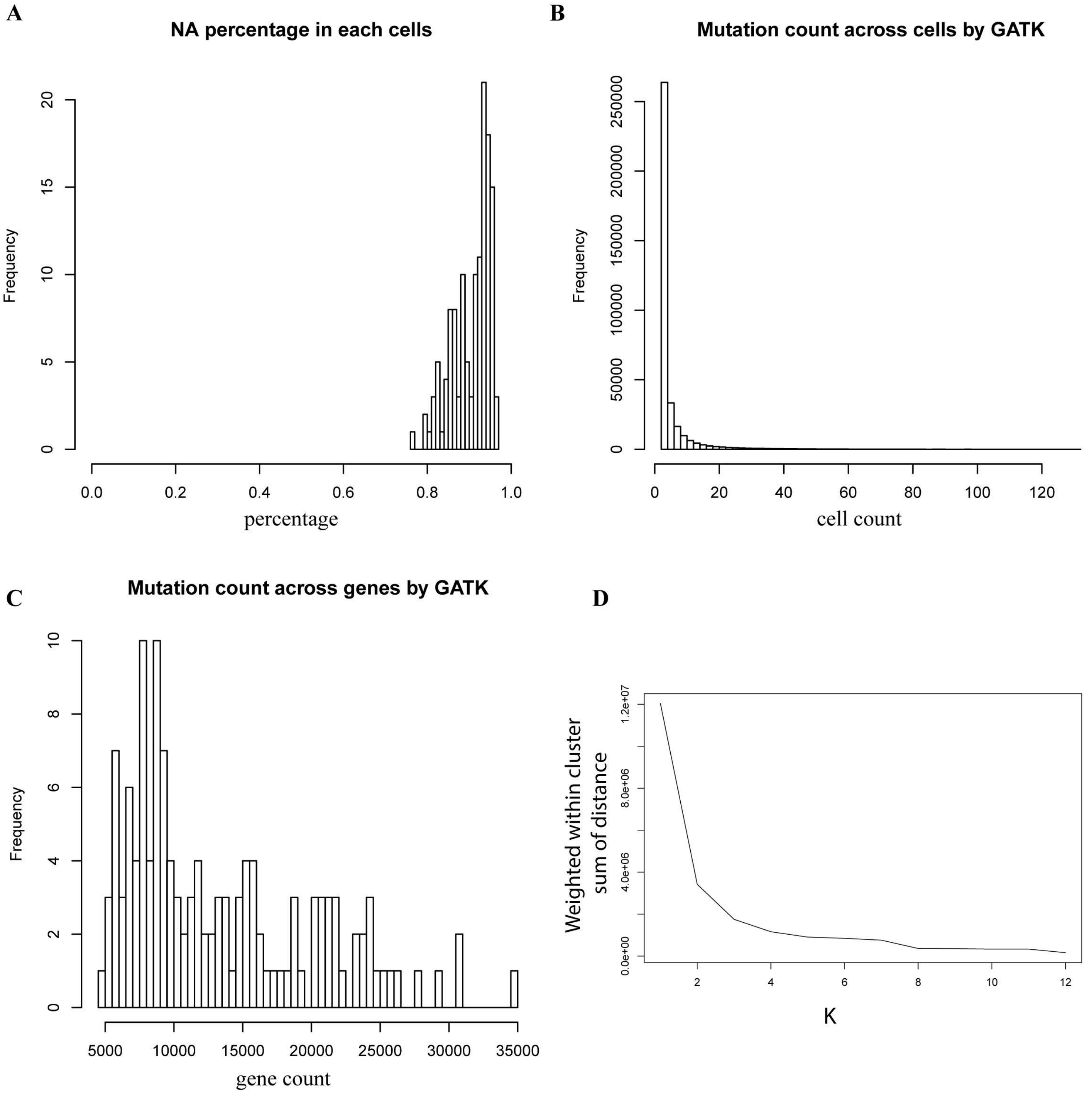


**Figure S12. Breast cancer dataset mutation statistics detected by GATK tool and optimal clustering option. a** NA percentage in each cell across genes. When there is no read counts, it shows as NA. **b** Mutation count across cells by GATK. Most of the genes have low mutation frequency. **c** Mutation count across genes by GATK tool. It shows mutation counts with a bell shape. **d** Intra-cluster divergence curve given different number of clusters (K). 5 is the elbow point.

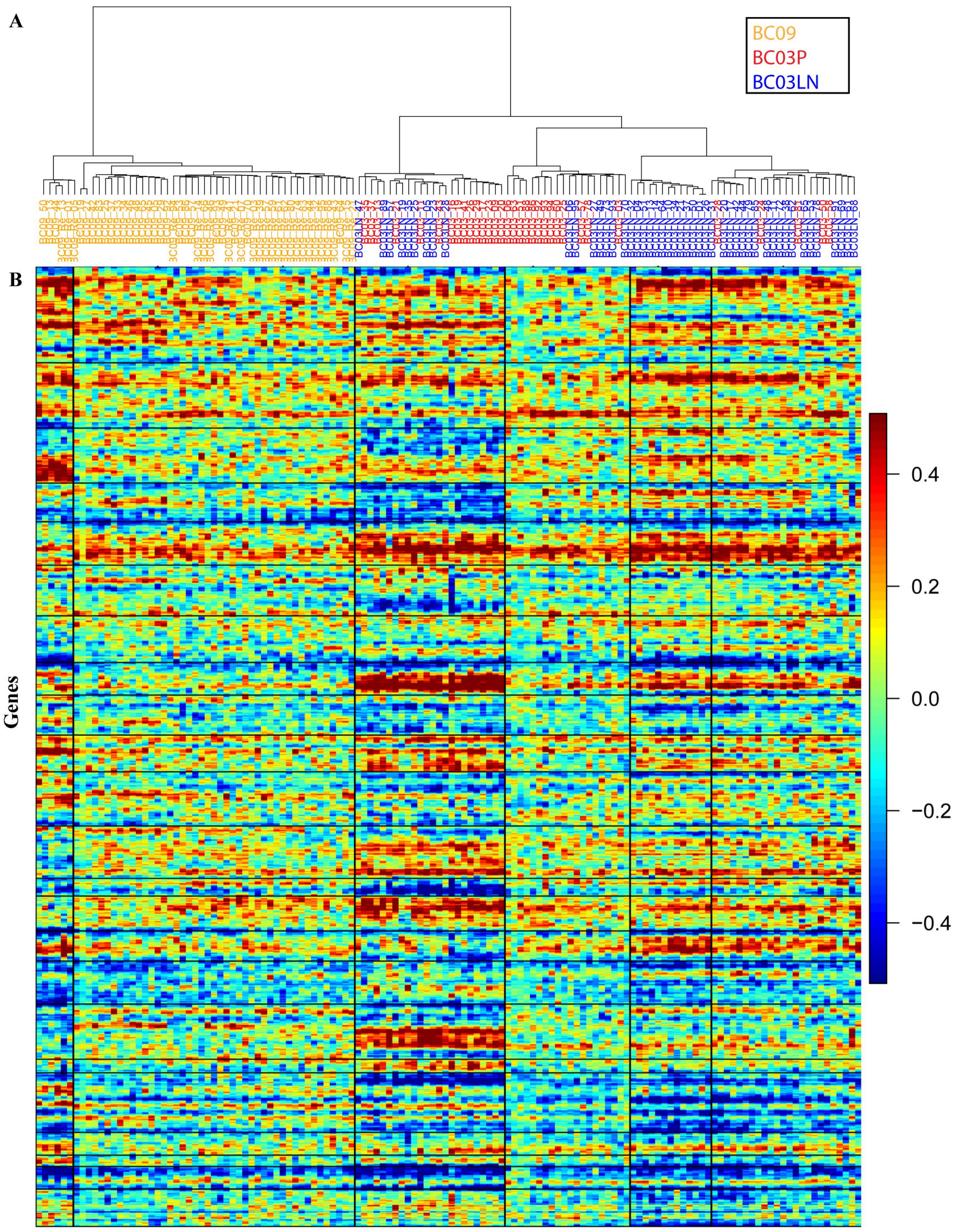


**Figure S13. Expression of primary breast cancer. a** DENDRO clustering of breast cancer. **b** Expression ordered by DENDRO clustering

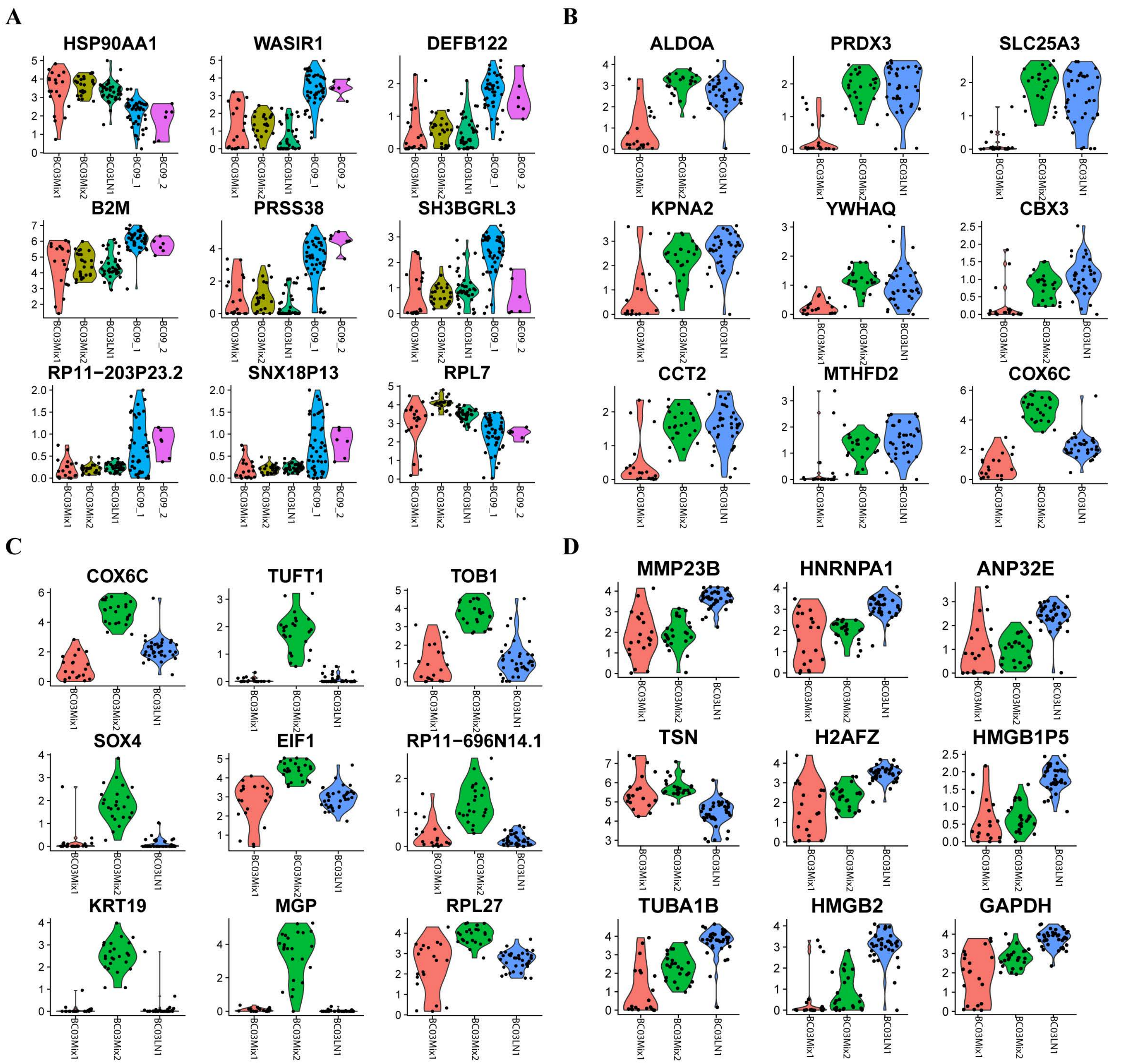


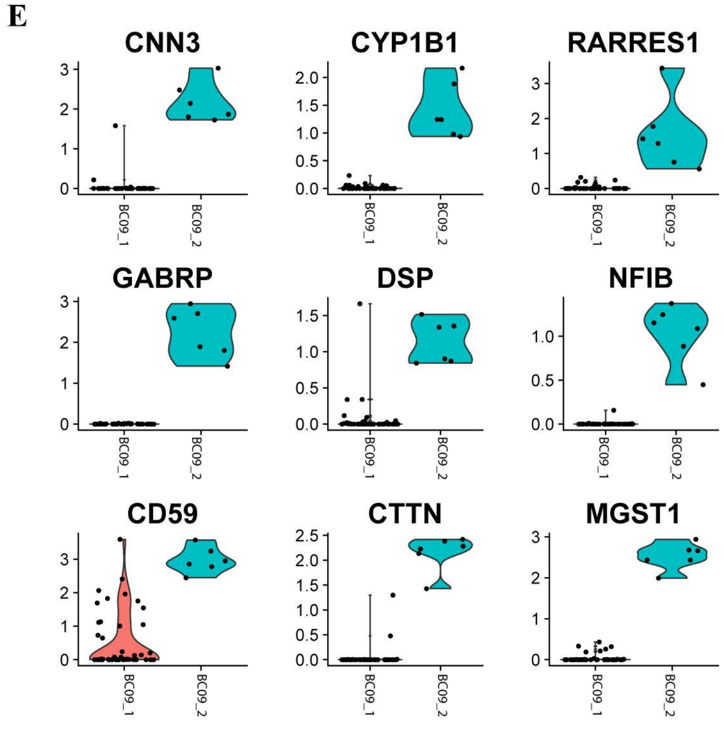


**Figure S14. Most significant differential expressed genes between different BC pairs. a** BC03 vs. BC09. **b** BC03Mix1 vs. BC03 other. **c** BC03Mix2 vs. BC03 other. **d** BC03LN1 vs. BC03 other. e BC09_1 vs. BC09_2
